# The human RNA polymerase I structure reveals an HMG-like transcription factor docking domain specific to metazoans

**DOI:** 10.1101/2021.12.22.473891

**Authors:** Julia L. Daiß, Michael Pilsl, Kristina Straub, Andrea Bleckmann, Mona Höcherl, Florian B. Heiss, Guillermo Abascal-Palacios, Ewan Ramsay, Katarina Tlučková, Jean-Clement Mars, Astrid Bruckmann, Carrie Bernecky, Valérie Lamour, Konstantin Panov, Alessandro Vannini, Tom Moss, Christoph Engel

**Affiliations:** Regensburg Center for Biochemistry, University of Regensburg, 93053 Regensburg, Germany; Division of Structural Biology, The Institute of Cancer Research, London SW7 3RP, United Kingdom; Institute of Science and Technology, Am Campus 1, 3400 Klosterneuburg, Austria; Department of Molecular Biology, Medical Biochemistry and Pathology, Faculty of Medicine, Laval University, Quebec, QC, G1V 0A6, Canada; Laboratory of Growth and Development, St-Patrick Research Group in Basic Oncology, Cancer Division of the Quebec University Hospital Research Centre, Québec, Canada; Université de Strasbourg, CNRS, INSERM, Institut de Génétique et de Biologie Moléculaire et Cellulaire (IGBMC), Department of Integrated Structural Biology, Illkirch; Hôpitaux Universitaires de Strasbourg, Strasbourg, France; School of Biological Sciences and PGJCCR, Queen’s University Belfast, Belfast, UK; Fondazione Human Technopole, Structural Biology Research Centre, 20157 Milan, Italy

## Abstract

Transcription of the ribosomal RNA precursor by RNA polymerase (Pol) I is a major determinant of cellular growth and dysregulation is observed in many cancer types. Here, we present the purification of human Pol I from cells carrying a genomic GFP-fusion on the largest subunit allowing the structural and functional analysis of the enzyme across species. In contrast to yeast, human Pol I carries a single-subunit stalk and *in vitro* transcription indicates a reduced proofreading activity. Determination of the human Pol I cryo-EM reconstruction in a close-to-native state rationalizes the effects of disease-associated mutations and uncovers an additional domain that is built into the sequence of Pol I subunit RPA1. This ‘dock II’ domain resembles a truncated HMG-box incapable of DNA-binding which may serve as a downstream-transcription factor binding platform in metazoans. Biochemical analysis and ChIP data indicate that Topoisomerase 2a can be recruited to Pol I via the domain and cooperates with the HMG-box domain containing factor UBF. These adaptations of the metazoan Pol I transcription system may allow efficient release of positive DNA supercoils accumulating downstream of the transcription bubble.

## Introduction

Transcription of DNA into RNA is carried out by three nuclear polymerases (Pols) in most higher eukaryotes^1^. These multi-subunit Pols diverge in target loci, structure and regulation ^2^. Understanding the underlying molecular mechanisms is a central goal of molecular biology. However, these mechanisms have been mostly studied in lower model organisms due to experimental limitations. In higher eukaryotes, regulatory variations dependent on tissue type, developmental state and cell-cycle stage are adding additional layers of complexity. The structure-function analysis of human Pol II^3^ and Pol III^4–7^ showed both similarities in the catalytic mechanisms and divergence in regulatory elements among organisms.

Human RNA polymerase (hPol) I has a single target gene, the 47S ribosomal RNA precursor (pre-rRNA), from which the 5.8S, 18S and 28S rRNA are processed^8^. These processed RNAs contribute to ribosome formation together with the 5S rRNA synthesized by Pol III^9^. rRNA synthesis contributes up to 80% of total cellular RNA^10^ and must therefore be tightly regulated. Hence, dysregulation of hPol I is associated with pathologies, such as cancer and developmental diseases, for example Treacher Collins Syndrome^11^. Unsurprisingly, inhibition of hPol I has been explored as a therapeutic strategy with some success in cancer treatment and future potential^12^. The molecular action of rRNA synthesis inhibitors is not entirely understood and may range from the activation of DNA-damage responses upon interference with replication^13^ to a specific reduction of Pol I transcription by preventing promoter escape during initiation^14^ or inhibiting elongation^15^.

The composition of hPol I is similar to yeast Pol I^16^ of which detailed crystal structures are known^17,18^. A catalytic core of ten subunits is complemented by a protruding stalk subcomplex and a heterodimeric RPA49/RPA34 subcomplex. The latter is related to Pol II initiation factors TFIIF and TFIIE^19^ and has homologues in Pol III^20^. The stalk was proposed to be divergent between yeast and human, as DNA- and protein sequence based searches have not identified an homologue of subunit A14 in human cells^16^. Table 1 summarizes the subunit terminologies for yeast and mammalian Pol I in comparison to human Pol II and Pol III subunits and correlates nomenclature. Regulation of Pol I is diverse^21^ and can be achieved by post-translational modification (PTM) of Pol I subunits or transcription factors. Nutrient availability^22^ and growth factor signal transduction^23^ activate Pol I initiation by phosphorylation of initiation factor Rrn3. Rrn3 is essentially conserved among species^24–26^ and primes Pol I for initiation by interacting with the stalk subcomplex^27–30^. Furthermore, dephosphorylation of the stalk is required for efficient Pol I function in yeast^31^ and hyper-acetylation of RPA49 reduces Pol I activity under stress^32^.

**Table 1.**
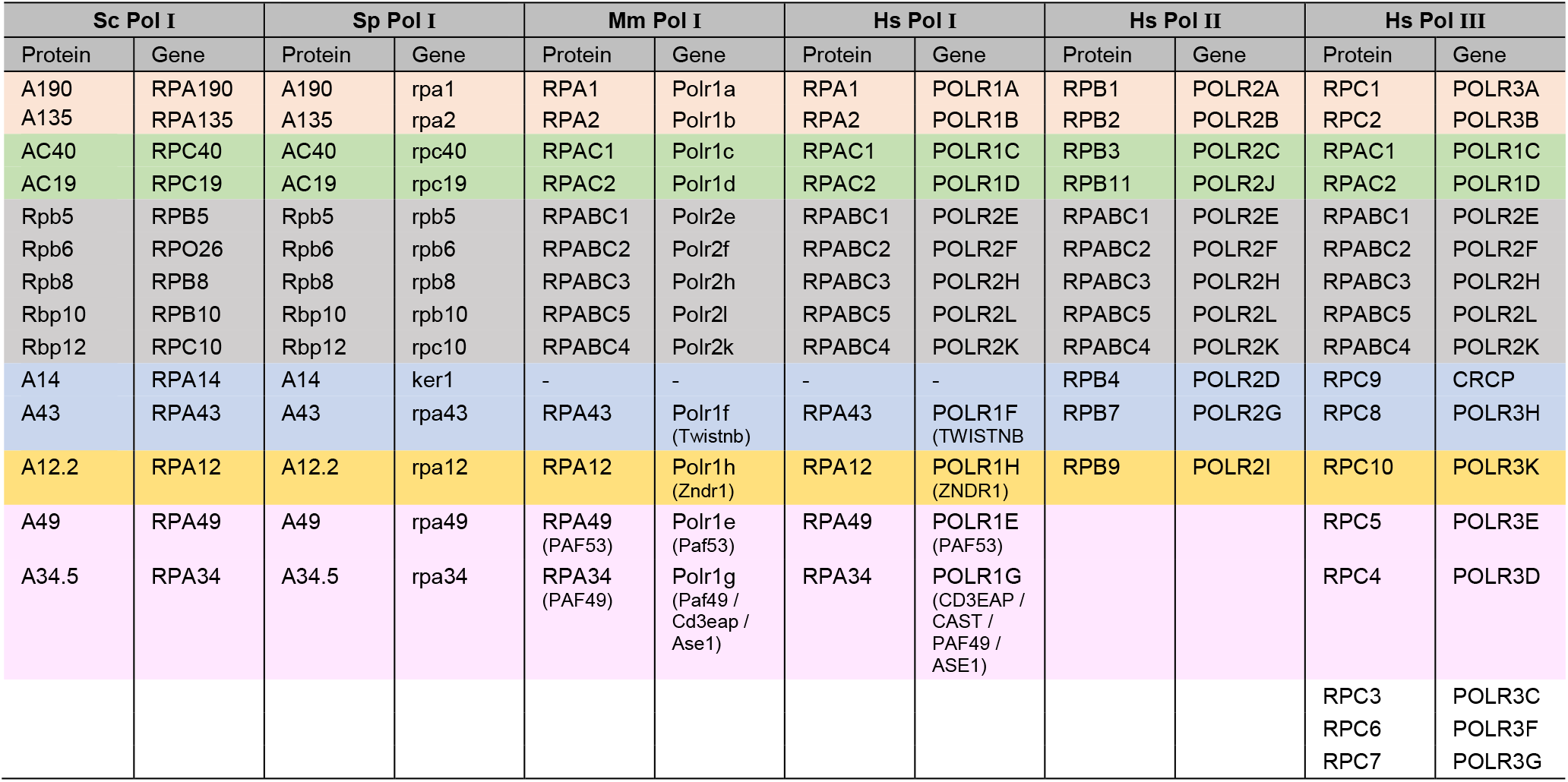
Human Pol I, II and III subunit nomenclature in relation to yeast counterparts. Wheat: Large subunits; Green: subunits shared between Pol I and III; Grey: Common subunits; Blue: Stalk sub-complex; Orange: Rpb9/TFIIS-like subunits; Pink: Built-in TFIIF/E-like subunits of Pol I and III.

Functionally, hPol I transcription has been studied in extracts or partially purified systems^33^. In contrast, yeast Pol I transcription could be studied in detail using purified and recombinantly expressed components, allowing a clear definition of subunit functionalities in transcription initiation^28,34^, elongation^35,36^, cleavage^37^, backtracking^38,39^ and termination^40,41^. Such studies allowed a detailed dissection of (sub-)domain and transcription factor functions.

Due to the lack of a well-defined *in vitro* system consisting of purified components, it is unclear whether the results of structure-function studies can be transferred to higher organisms. Apparently, many factors are conserved functionally but diverge in composition^42^. In addition to RRN3 the hPol I transcription requires the initiation factors ‘Selectivity Factor 1’ (SL1) and ‘upstream binding factor’ (UBF). SL1 comprises the subunits TAF1A, TAF1B, TAF1C (homologues of yeast Core Factor), the two additional factors TAF1D^43^ and TAF12^44^, and includes the TATA-binding protein (TBP). UBF consists of six consecutive HMG boxes, is a part of initiation complexes^45^ and binds to the body of actively transcribed rDNA genes^46^, apparently preventing re-association of nucleosomes.

It remains poorly understood how Pol I structurally and functionally adapted to the increased regulatory demands in human cells. Here, we show how hPol I can be exclusively purified from a modified human cell line in its natural form and determine its structure by single-particle electron cryo-microscopy (cryo-EM). The structure reveals a previously unknown, built-in platform that may allow docking of transcription factors on the downstream face of the polymerase. Phylogenetic analysis allows following the evolution of Pol I by the loss of a subunit and the gain of additional domains in higher organisms. We present *in vitro* transcription assays demonstrating a limited proofreading ability of the human enzyme and map known mutations on the structure to understand Pol I -related pathologies.

## Results

### Specific tagging and purification of human RNA polymerase I

To study the structure and function of hPol I *in vitro*, we first created a cell line that allows the specific enrichment of the complete enzyme in its native state without contamination of hPol III. Using the CRISPR/Cas9 technology in a dual-nicking approach, a cleavable sfGFP tag was fused to the genomic sequence of the largest Pol I subunit RPA1 of the Hela P2 cell line^47^. Following identification of positive clones by single-cell FACS based on GFP fluorescence intensity, correct insertion was confirmed by site-specific PCR. Homozygous insertion was verified by western blot against subunit RPA1 (Fig. 1). The approach we previously reported for the generation of an RPAC1-tagged cell line^4^ can hence be generally applied for reliable homozygous knock-in of C-terminal fusion tags.

**Figure 1.**
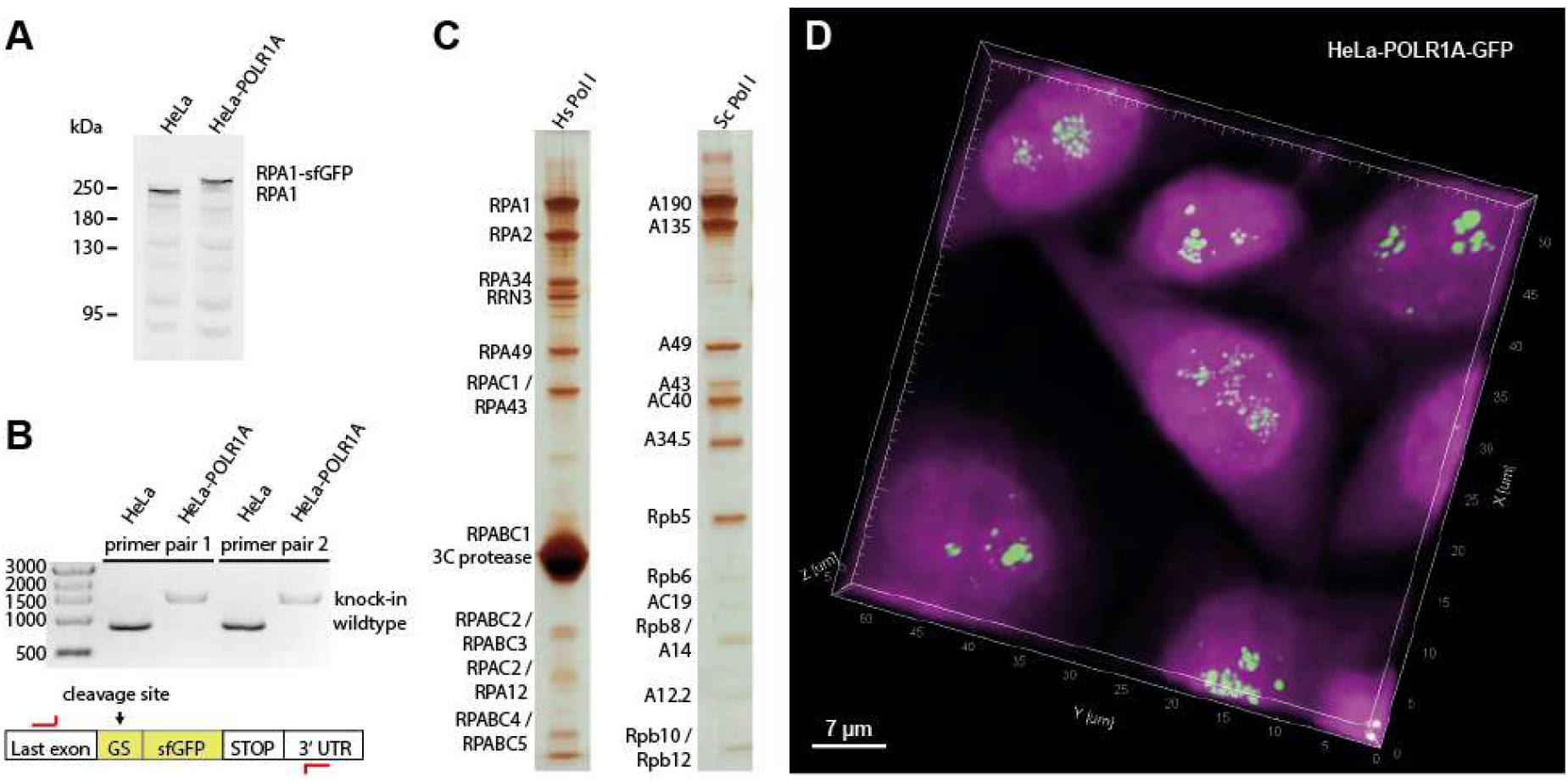
Homozygous sfGFP knock-in cell line generation and human Pol I purification. **A** Western blot against RPA1 shows a shift to larger molecular weight in lysates of the RPA1-sfGFP cell line, confirming exclusive expression of the modified protein. **B** Site-specific knock-in of the cleavable sfGPF fusion confirmed by PCR from genomic DNA (Sybr-Safe stained agarose gel). **C** Purification of human Pol I shows bands for all subunits in comparison to the *S. cerevisiae* enzyme (silver-stained SDS-PAGE). **D** Confocal imaging shows the exclusive location of GFP-induced fluorescence in the nucleoli in aligned 3D stacks. Spots in the central cell may represent single rDNA genes. Magenta: DAPI stain; Green: sfGFP signal (fused to RPA1).

hPol I purification from lysates of the RPA1-sfGFP cell line relies on a single affinity purification step followed by site-specific tag-cleavage, resulting in a highly enriched sample (Fig. 1C; Sup. Fig. 1). As judged by mass spectrometry (Sup. Fig. 2), the sample partially co-purifies with the initiation factor RRN3 and contains stoichiometric amounts of hPol I subunits, including the RPA49/RPA34 sub-complex, which is sub-stoichiometric in rat Pol I purifications^48^. An optional subsequent ion-exchange chromatography step resulted in the loss of initiation factor RRN3 and the RPA49/34 subcomplex from most polymerases (Supplemental Fig. 1B).

### Human Pol I shows reduced proofreading *in vitro*

Equipped with a cell line that allows the specific enrichment of hPol I, we now aimed at a detailed structural and functional characterization of this enzyme *in vitro*. To understand functional conservation, we first compared purified hPol I activity with its counterparts from *S. cerevisiae* and *S. pombe* in an *in vitro* elongation and cleavage assay. A fluorescently labeled RNA primer is extended in the presence of nucleotide triphosphates (NTPs) by Pol I, or cleaved due to the action of the TFIIS-related subunit RPA12 (Fig. 2A). While yeast Pol I specifically incorporates the correctly base-paired substrate, hPol I generates transcripts containing incorrectly incorporated NTPs under identical experimental conditions (Fig. 2B).

**Figure 2.**
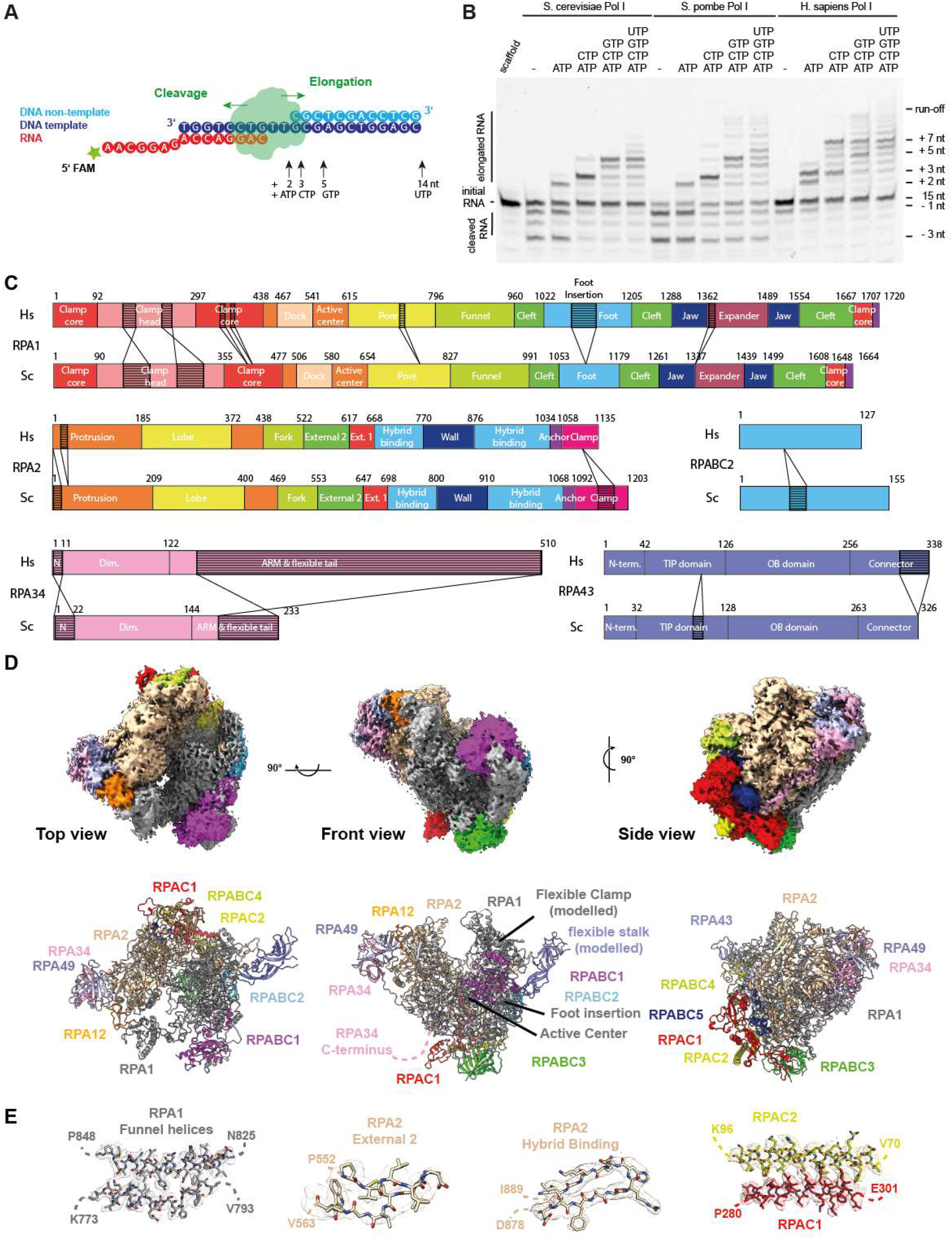
Activity, domain architecture and cryo-EM reconstruction of human Pol I. **A** Schematic representation of the assays and scaffold sequences used to determine hPol I activity *in vitro*. **B** Compared to Sc and Sp Pol I, cleavage activity of hPol I is reduced and only reaches the -1 position, while Sc and Sp enzymes can cleave up to three nucleotides from a matched hybrid. Elongation efficiency is comparable, although incorporation of mis-matched nucleotides is strongly increased in the case of hPol. **C** Schematic domain architecture of the Pol I subunits with largest differences to their yeast Pol I counterparts: RPA1, RPA2, RPA34, RPA43 and RPABC2. Subdomains and insertions/deletions of ten or more residues indicated. **D** Cryo-EM density of human Pol I shows flexibilities in the clamp/stalk region of RPA1 and RPA43. Structure model shown below. **E** Enlarged view of RPA1 funnel helices, RPA2 External II and Hybrid Binding domains and the RPAC1/2 assembly overlaid with sharpened cryo-EM density.

Furthermore, the cleavage pattern of yeast and human Pol I in the absence of NTPs diverges. While the 3’-end of the perfectly base-paired RNA primer can be cleaved up to three nucleotides by Sc and Sp Pol I, the main product of hPol I cleavage is at position -1, indicating a reduced backtracking ability. To exclude effects from potential sub-stoichiometry of the RPA49/34 complex, we added recombinantly co-expressed human RPA49/34, but observed neither increased backtracking/cleavage, nor reduced generation of mismatched transcripts (Sup. Fig. 3B). Similarly, the addition of recombinant Rrn3 to Sc Pol I does not hamper is functionality (Sup. Fig. 3D), suggesting that the observed effects do not originate from RRN3 present in the sample.

To test the influence of the substrate scaffold, we added a non-template (nt) strand with a mismatched bubble and tested a wealth of different template sequences (Sup. Fig. 3 D-I). On a mismatched bubble-template, backtracking is impaired even further, while the incorporation of incorrect NTPs generally remained, but showed some sequence specific variations in intensity. Functional analysis of substrate mis-incorporation rates indicated a similar effect when Sc Pol I is compared to Sc Pol II^49^. This is well in line with our observations and may originate from the flexibility among Pol I core and shelf modules as discussed^50,51^. To understand the evolution of Pol I and to rationalize the functional differences between the enzymes of different species, we determined the structure of human Pol I by cryo-EM.

### Structure determination of hPol I

Whereas the structure of yeast Pol I has been extensively studied by X-ray crystallography^17,18,52^ and single-particle cryo-EM^53–55^, the human enzyme eluded structural characterization thus far. In a first step, negative stain EM screening revealed intact particles (Sup. Fig. 1C-E) and a 3D reconstructed negative stain envelope indicated an architecture comparable to *S. cerevisiae* Pol I. However, many particles show flexibilities in the clamp/stalk region that originate from heterogeneity or functional flexibility. High-resolution structure determination by single-particle cryo-EM was hampered by intrinsic flexibility and a strong bias in orientation distribution of hPol I particles. Finally, data collected from self-made graphene oxide-covered grids reduced orientational bias of non-crosslinked particles after extensive screening for preparation conditions (Pilsl et al., Methods Mol Biol, in press). We collected a total of 9,709 micrograph movies on a CryoARM 200 (JEOL) electron microscope equipped with K2 direct electron detector (Gatan) at a pixel size of 0.968Å. Preprocessing and particle picking in Warp^56^ was followed by binning and 2D classification in RELION 4.0^57^, yielding 145,554 particles that were subsequently subjected to sequential 3D classification (Sup. Fig. 4; Sup. Table 1). A 3D reconstruction with an overall resolution of 4.09 was obtained, revealing secondary structures for most regions of the molecule. Models for common subunits RPABC1-ABC5 and the RPAC1/2 assembly were transferred from a hPol III reconstruction^5^. Homology models of the hPol I subunits RPA1, RPA2, RPA49, RPA34, RPA12 and RPA43 were generated based on sequence and secondary structure alignments with the crystal structures of their *S. cerevisiae* counterparts (Sup. Data 1) using the MODELLER software package^58^. Model fitting and rigid body refinement allowed interpretation of both negative stain and cryo-EM densities and later aided by AlphaFold predictions^59^.

To the knowledge of the authors, this is the first example for the *de novo* reconstruction of a previously unknown, non-symmetric macromolecule obtained with a CryoARM 200 electron microscope. Details of data collection and handling strategies are similar to recent reports^60–62^ and are described in the methods section.

### Insights into hPol I architecture

The negative stain density shows both, the stalk and the RPA49/RPA34 heterodimer (Sup. Fig. 1E). However, some 3D classes lack density for the region of the clamp core and clamp head domain of subunit RPA1 and the stalk, indicating a high flexibility of this sub-assembly.

The cryo-EM reconstruction of hPol I (Fig. 2D) shows connected density for the common hPol subunits RPABC1-5, the RPAC1/2 dimer, the N-terminal domain of subunit RPA12 and most parts of subunit RPA2, with exception of the C-terminal clamp and anchor domains (residues 1010-1134). Furthermore, density for the jaw, funnel, foot and most parts of the cleft domain of subunit RPA1 (residues 630-1661 excluding loops) and for the RPA49/34 heterodimer allowed unambiguous fitting of homology models. In our reconstruction, weak density for the stalk subcomplex, the clamp and dock domains of subunit RPA1 indicate increased shelf module flexibility. Similar to yeast Pol I crystal structures, the linker and tWH domains of subunit RPA49 and the C-terminal extension of subunit RPA34 are also flexible in human Pol I.

The assembly of RPAC1/2 reflects the conformation known in hPol III and tightly interacts with subunit RPA2. The N-terminus of subunit RPA12 can be placed on the lobe of subunit RPA2, demonstrating the stable association of the subunit. Global contraction of Pol I modules upon activation has been observed in the enzymes of *S. cerevisiae*^17,18^ and *S. pombe*^63^ and may be a regulatory feature of Pol I^50,51^. Negative stain EM and cryo-EM sample freezing of hPol I complexes without the use of crosslinking reagents to artificially stabilize conformations may indicate a close-to-native state of functional importance in the human Pol I. Overall, the architecture of hPol I reflects that of the yeast counterparts, but allows insights into the effects of Pol I-related mutations identified in human disease and reveals two major adaptations accumulating upon evolution: the stalk sub-complex (flexible in our density) and the RPA1 foot domain.

### Mapping of disease-associated mutations to Pol I subunit structures rationalizes enzyme deficiencies

Four disease phenotypes were linked to mutation of Pol I subunits in humans: Acrofacial Dysostosis (Cincinnati type)^64,65^, Treacher-Collins syndrome (TCS)^66–69^, Hypomyelinating Leukodystrophy (HL)^68,70^, and a juvenile neurodegenerative phenotype akin to the HL-phenotype^71^. With the structural model of hPol I determined (Fig. 2), we mapped these known mutations to gain insight into the underlying molecular pathologies (Sup. Fig. 6).

Acrofacial Dysostosis, Cincinnati type, leads to craniofacial abnormalities during development and is caused by mutations E593Q and V1299F in subunit RPA1^64,65^. Mutation E593Q is located in proximity to the catalytic center and may directly affect the nucleotide addition (Sup. Fig. 6c). In contrast, V1299F is situated on the interface of RPA1 with RPA12 and may destabilize the association of this subunit with the hPol I core (Sup. Fig. 6d).

Treacher-Collins syndrome (TCS) is a craniofacial developmental disease caused by various mutations in the genes TCOF1, POLR1B, POLR1C or POLR1D. Serine 682 of RPA2 directly contacts the bridge helix (likely H967 of RPA1) which may be affected by the mutation resulting in partially hindered translocation (Sup. Fig. 6f). In contrast, R1003 of subunit RPA2 is situated in the DNA/RNA binding cleft and may be required to stabilize folding of the hybrid-binding domain within RPA2 (Sup. Fig. 6g). Hence R1003C and R1003S^69^ may lead to a destabilization of RPA2 and thus the active center. Other TCS-associated mutations within subunit RPAC2 (E47K, T50I, L51R, G52E, L55V, R56C, L82S, G99S) cluster at intra-subunit and RPAC1 inter-subunit contacts^66–68^ (Sup. Fig. 6h). Structural alignment with the human Pol III structure reveals a similar fold and suggests destabilizing effects of these mutations, similar to R279Q/W of subunit RPAC1. Therefore, polymerase-associated TCS mutations can be functionally classified according to their effects: (1) Impaired Pol I transcription activity (RPA2 mutations) and (2) Effect on Pol I and Pol III transcription.

Similar to TCS, Hypomyelinating Leukodystrophy (HL) is a neurodegenerative disease that cannot be classified as a Pol I- or Pol III-associated disease per se. HL mutations are found in subunit RPAC1 which is shared between both polymerases or in Pol III subunits RPC1 and RPC2^68,70^. Comparing the structures of hPol I and hPol III shows that mutations of the RPAC1 N-terminus (T26I, T27A, P30S, N32I) are likely to have a Pol III-specific effect as this region appears flexible in hPol I, but mediates interactions to the polymerase core in Pol III (Sup. Fig. 6i). The nearby mutation N74S (as N32I) affects Pol III assembly but apparently does not impair Pol I biogenesis or nuclear import^68^. Additionally, RPAC1 mutations I105F, H108Y and R109H were found to impair RPC2 interaction in hPol III but not RPA2 in Pol I, again suggesting Pol III-specificity (Sup. Fig. 6j). Additional RPAC1 mutations M65V, V94A, A117P, G132D, C146R, R191Q, I262T, T313M and E324K are involved in the formation of intra-subunit contacts, likely affecting RPAC1 folding itself (Sup. Fig. 6e).

Finally, the mutation S934L in RPA1 is associated with a juvenile neurodegenerative phenotype akin to the HL-phenotype associated with Pol III disruption^71^. This mutation occurs in a small loop of RPA1 which forms contacts with RPA2 in the vicinity of the bridge helix N-terminus (Sup. Fig. 6b). This may generally disrupt and destabilize the Pol I core to some extent.

### A single-subunit stalk is the predominant configuration for Pol I

One of the major differences between Pol I enzymes of different organisms lies within the stalk subcomplex. DNA- and protein-sequence based searches identified homologues for 13 of the 14 yeast Pol I subunits except for the stalk-subunit A14 ^16^. Divergence of the stalk subunits among DNA-dependent RNA polymerases is well documented. Compared to the Pol II stalk, a domain-swap between yeast Rpb4 and Rpb7 and the yeast Pol I stalk subunits A14 and A43 was observed in the crystal structure of the Pol I subcomplex^37,72^. With this swap, subunit A14 appears to harbor limited functional importance. Deletion of the subunit in *S. cerevisiae* is not lethal but results in conditional growth defects indicating regulation deficiencies^73,74^, similar to observations in *S. pombe*^75^.

To analyze whether hPol I indeed carries a single-subunit stalk, mass spectrometric analysis of all protein bands in our purification was performed. The 13 subunits identified *in situ* and initiation factor RRN3 were found to be present with sequence coverages over 25 % (Sup. Fig. 2). Additional proteins were not identified with similar confidence. To clarify whether the absence of a second Pol I stalk subunit is specific to human cells and to understand the changed composition of the enzyme during its evolution, we carried out bioinformatic analysis: First, we generated a phylogenetic tree based on sequence similarity of the Pol I subunits RPA1, RPA34 and RPA43 to cover the polymerase core and the peripheral sub-complexes (Fig. 3). Generating a Pol I-specific conservation tree removed bias that may originate from the influence of unrelated genes on global alignments in standard phylogenetic analysis. We clearly find that only organisms of the *Saccharomycotina* in the *Dikarya* clade carry sequences for the subunit A14, indicating that a single-subunit stalk is the standard Pol I configuration.

**Figure 3.**
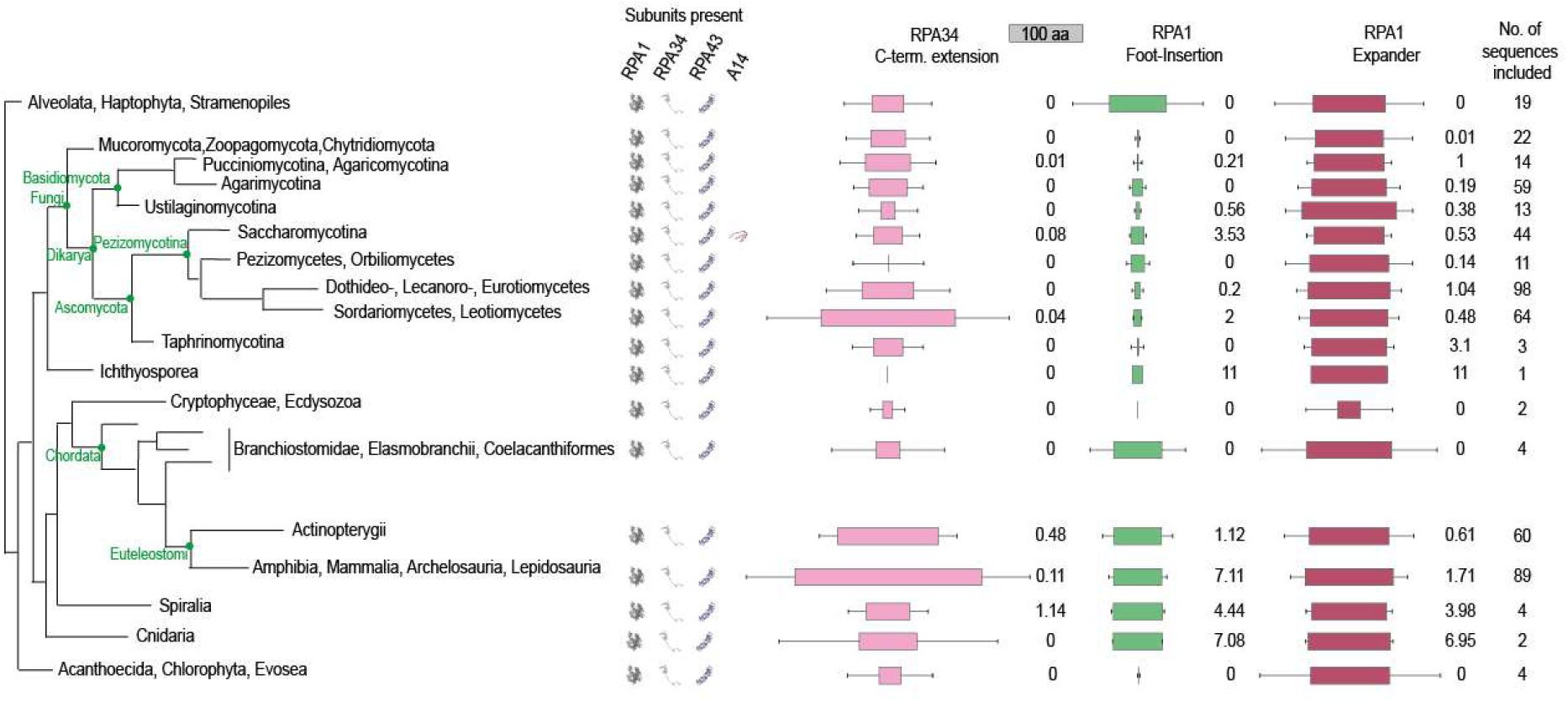
Phylogenetic analysis of RNA polymerase I. Phylogenetic tree calculated based on sequence homology of the three Pol I subunits RPA1 (core), RPA34 (RPA49/34 heterodimer) and RPA43 (stalk subcomplex); schematic. The subunit A14 is found in all *Saccharomycotina* in the class of *Dikarya*. This includes model organisms such as *S. cerevisiae* and *S. pombe*, explaining the current paradigm that Pol I comprises 14 subunits. Conservation scores for the RPA1 foot insertion, the Expander (DNA-mimicking loop) and the C-terminal extension of RPA34 were calculated in each class. Blocks show the median length of each specific region (100 residue referenced above). Box reflects the median; error bars indicate standard deviation; conservation scores are grouped into five categories: not conserved (0-3), weakly conserved (3-5), medium conserved (5-7), conserved (7-9) and strongly conserved (9-11).

### Built-in transcription factors differ among organisms

Phylogenetic analysis also showed that the ‘expander’ (DNA-mimicking) element is present in all analyzed organisms. This flexible insertion in the jaw domain of the largest subunit mimics DNA binding to inactive Pol I dimers^17,18^ or monomers^63^.

The RPA49/34 heterodimer resembles the yeast A49/A34.5 sub-complex with functions in initiation and elongation^28,35,76^ and is present in cryo-EM reconstructions. The subcomplex is related to the Pol II initiation factors TFIIF and TFIIE^19^ and stays attached to the Pol I core throughout its transcription cycle *in vivo*^77^, but may be lost under some conditions *in vitro*^37,55,63^. The TFIIE-related, C-terminal tWH domain of subunit RPA49 is flexible in our reconstructions as expected for Pol I monomers and most elongation states. Similarly, we do not observe density for the mammalian-specific C-terminal extension of subunit RPA34 (compare Fig. 2/3). This is also the case for a C-terminal extension of the hPol III subunit RPC5 that contributes to enzyme stability despite being flexibly linked^4^.

The C-terminal domain of RPA34 is enlarged to 55 kDa in humans compared to the 27 kDa yeast protein (Fig. 2C; Supplementary Data 1). The C-terminal extension is present in higher organism classes, such as *Mammalia* and *Amphibia*, but shows no clear conservation in sequence, predicted secondary structure or length (Fig. 3), and is flexible in our cryo-EM reconstruction. To determine functional similarity with the yeast counterparts, we tested binding of recombinant human RPA49/34 to the *S. cerevisiae* enzyme purified from an A49 deletion strain resulting in a 12-subunit Pol I (Pol IΔ). Direct cross-species binding of the RPA49/34 heterodimer to Sc Pol I *in vitro* was not possible, likely due to divergence of the charged tail region (‘ARM’) of RPA34 and its binding site on the ‘external’ domain of the second largest subunit RPA2.

In contrast to direct interaction, functional cross-species complementation of recombinant yeast and human sub-complexes was possible (Sup. Fig. 3C). Recombinant Sc A49/34.5 and Hs RPA49/34 both recovered the activity of hPol IΔ in elongation and cleavage. Hence, interaction interfaces apparently co-evolved, while subcomplex function was retained from yeast to human. Both, Sc and Hs RPA49/34 can bind to DNA independent of core Pol I (Sup. Fig. 5C). While the main interface with DNA apparently lies within the TFIIE-related tWH domain of RPA49, the flexible and divergent RPA34 tail is capable of independent DNA-interaction. Notably, the elongation and cleavage pattern indicated no major differences depending on the type of heterodimer added (Sc or Hs version). Therefore, reduced proofreading of hPol I apparently is an intrinsic enzymatic feature of the core enzyme rather than effects introduced by divergent heterodimer subunits or their sub-stoichiometric co-purification.

### A previously undescribed domain is built into the largest subunit of human Pol I

The second major difference between yeast and human Pol I is an insertion in the ‘foot’ domain of the largest subunit RPA1 (Fig. 2C; Supplemental Data 1). The Pol II foot domain serves as transient interaction platform for the regulatory co-activator complex ‘mediator’^78^ and is enlarged compared to yeast Pol I^17,18^. This may lead to a speculation about a comparable regulatory role of the foot insertion specifically required in humans but not in yeast. We found well-defined cryo-EM density on the downstream face (front) of hPol I subunit RPABC1 (Rpb5) that is closely connected to the foot insertion site. Domain prediction using the HHPRED package^79^ indicated a clear homology to a High Mobility Group (‘HMG’) box domain with the closest fit to the structure of HMG box 5 of the hPol I transcription factor UBF^80^. Hence, we constructed a homology model of the foot insertion and fitted the resulting model into the observed cryo-EM density. This allows an unambiguous placement of the domain without adjustment, indicating that the hPol I foot insertion indeed resembles a built-in HMG box (Fig. 4).

**Figure 4.**
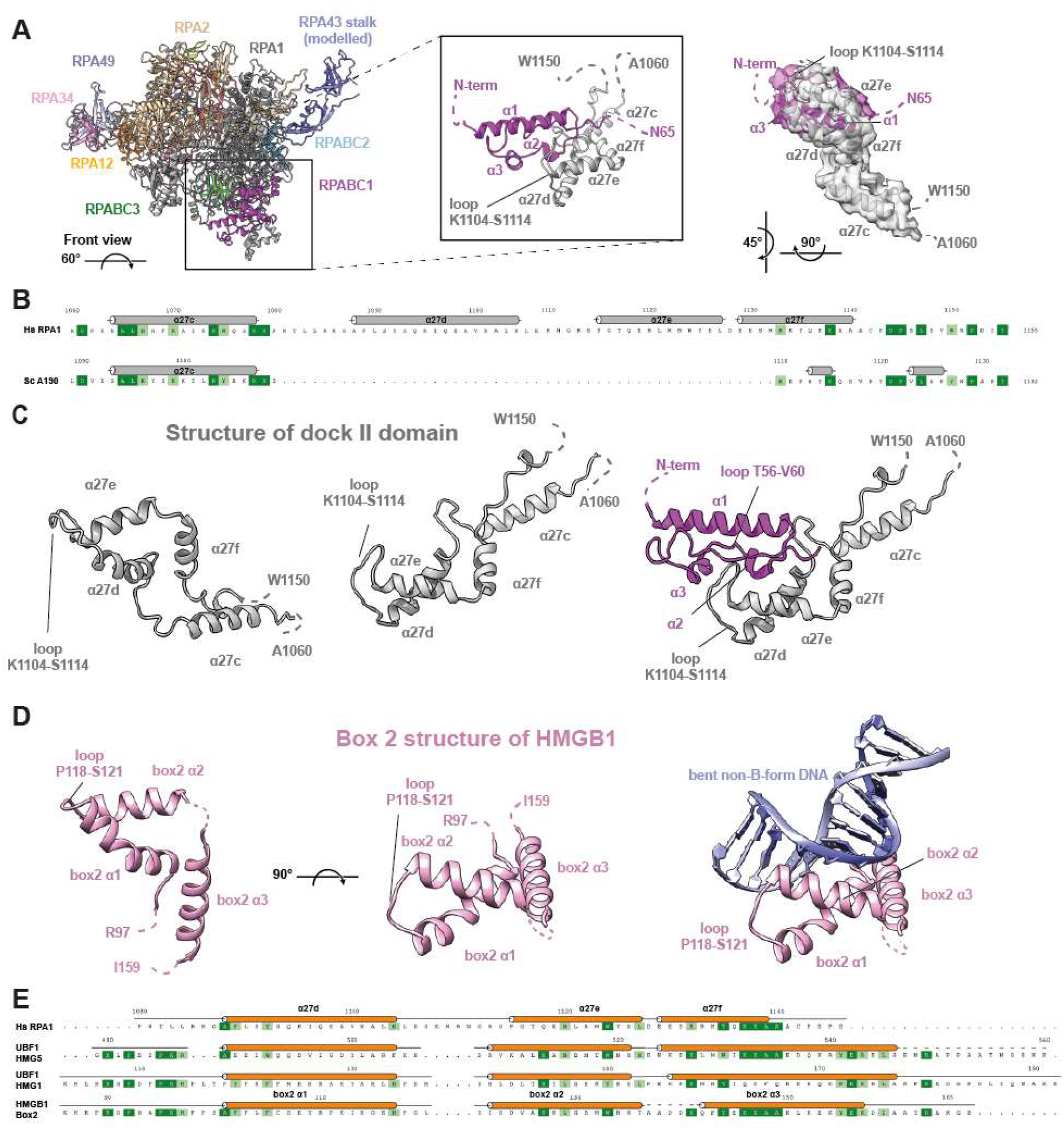
An HMG-box like domain is included into the largest subunit of human Pol I. **A** Location of the structured insertion in RPA1 α27d-f on the downstream edge of subunit RPABC1 in human Pol I and enlarged view of the region. Overlaid experimental cryo-EM density for the helices α27c-f of subunit RPA1 (grey) and the N-terminal 65 residues of RPABC1 (purple) shown as transparent surface (right). **B** Structure-based sequence alignment of human and yeast Pol I foot insertions (for complete sequence, compare Sup. Data 1). **C-D** Structure of the RPA1-foot insertion (**C**) compared to the canonical HMG box 2 of the human protein HMGB1 (**D**) from two views. The DNA-binding surface of the canonical HMG box 2 is occluded by RPABC1 in hPol I. **E** Structure-based sequence alignment of the RPA1-HMG insertion with the canonical HMG box 2 of HMGB1, and the boxes 1 and 5 of the Pol I transcription factor UBF. In the RPA1-HMG box, the N-terminal region is divergent and the third helix is truncated. Both of these parts are important for DNA-interaction. A loop insertion between the first two helices is part of the RPABC1-interface.

### The HMG box-containing ‘dock II’ domain may serve as interface for Topoisomerase 2a

Canonical HMG box domains can bind the minor groove of a DNA duplex in a sequence-specific or unspecific manner with a preference for non-B-form conformations^81^. Overlay with a model HMG box (box 2 of the human HMGB1 protein) shows that the DNA-binding site of the hPol I foot insertion is completely occluded by the common Pol subunit RPABC1 (Fig. 4C/D), indicating a divergent function. Furthermore, structure-based sequence alignment of the RPA1 foot HMG box shows that the so-called ‘minor wing’ is absent. This minor wing consists of an N-terminal motif and the C-terminal extension of the HMG helix three (Fig. 4E). Both regions cooperate in DNA-binding of canonical HMG boxes but are absent in RPA1. Furthermore, a loop between HMG-box helices one and two directly interacts with DNA and contributes to sequence specificity^82^. In RPA1, we observed an insertion between the corresponding helices α27d and α27e that contacts loop T56-V60 of subunit RPABC1 (Fig. 4). In contrast, a basic surface patch is found on the opposite face (Sup. Fig. 7). To test whether DNA-interaction is possible, we recombinantly expressed MBP-tagged versions of the domain (full length and minimal) and tested their ability to bind an unspecific dsDNA-fragment. Significant DNA-binding was not observed, although the full-length fragment may retain some very low affinity *in vitro*. We conclude that the RPA1 foot insertion represents a truncated HMG-box ‘major wing’ unable to bind DNA.

Apart from binding DNA, HMG boxes can promote interaction between proteins. This appears the most likely function for the RPA1 foot HMG-box, which we hence termed ‘dock II’. The human HMGB1 protein was found to interact with Topoisomerase (Top) 2a independent of DNA, while promoting the activity of this enzyme^83^. In fact, active Top2a co-purifies with the hPol I-RRN3 complex^84^ and was described to be part of the hPol I transcription initiation machinery^85^. Therefore, we asked whether recombinant human Top2a lacking the unstructured C-terminal domain^86^ can interact with the RPA1 dock II domain. Indeed, we observe a shift in native PAGE of full length, but not minimal dock II or the MBP-tag alone, indicating the possibility for transient interaction (Sup. Fig. 7E).

Consequently, we asked whether Top2a binding mapped to the rDNA gene in cells and whether it would hint towards a typical initiation factor behavior^85^. To this end, we re-analyzed previously published Top2a ChIP-Seq data from mouse cells^87^ and mapped the initiation factor TAF1B (part of SL1 and homologous to TFIIB^88,89^), UBF, Pol I^46^ and Top2a to the rDNA gene as described^90^. As shown in Fig. 5A, TAF1B maps to clear peaks at the spacer promoter and the main rDNA promoter, defining the transcription start site (TSS). Pol I is distributed over the gene body and the spacer promoter, as expected in growing cells. Strikingly, Top2a maps to the rDNA locus but does not show the profile of a classical initiation factor, such as RRN3 which peaks at the promoter and tails out in the 5’ region of the rDNA gene^46^. Instead, Top2a is present over the entire gene, with some peaks in the 3’ region. These peaks apparently overlay with the UBF-binding sites.

**Figure 5.**
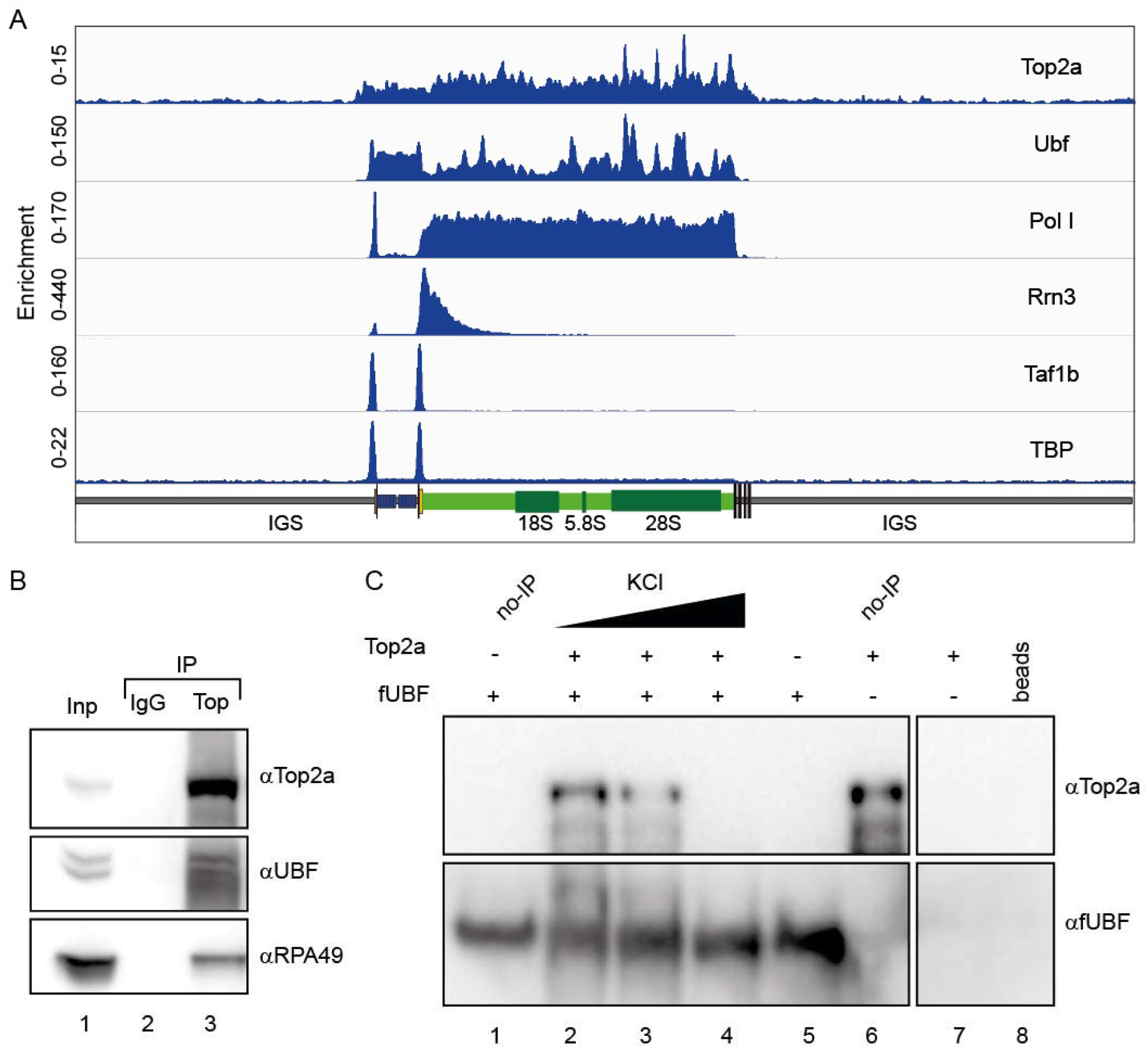
Top2a localizes to the rDNA gene and interacts with UBF. **A** Top2a is detected over the entire mouse rDNA gene regions occupied by UBF. Original raw data from^87^ was aligned and deconvoluted as previously described^90^. Peaks over the 3’ region of the gene overlap with UBF peaks, indicating co-localization. Top2a overlaps binding peaks for the initiation factors RRN3, TAF1B and TPB but specific correlations are not observed. Pol I signal marks the actively transcribed region. **B** UBF co-immunoprecipitates with Top2a: Top2a was immuno-precipitated from nuclear extract of U2OS cells using anti-Top2a antibodies (Abcam) immobilized on magnetic beads (DynaI). Immunoprecipitated proteins were analyzed by western blot using anti-UBF, anti-RPA49 and anti-Top2a antibodies Lane 1 – 10% input; Lane 2 – IP with IgG control; Lane 3 – IP with anti-Top2a antibodies. **C** Purified Top2a co-precipitates with purified UBF at low salt concentrations. Recombinant fUBF was incubated with purified Top2a at three different salt concentrations (lanes 2-4). Lane 1: fUBF control (no IP); Lane 5: IP without Top2a; Lane 6: Top2a control (no IP); Lane 7: IP without fUBF addition; Lane 8: FLAG-bead only control

### Physical interaction with UBF indicates functional cooperativity of Top2a and HMG-box containing proteins

Results from ChIP-Seq reanalysis do not exclude the possibility that Top2a is also part of some initiation complexes, but indicate either a Pol I - independent rDNA gene association, an elongation factor like behavior in cooperation with Pol I, and/or DNA-binding cooperativity with UBF. To test whether a physical interaction between UBF and Top2a takes place as indicated by co-localization of ChIP peaks, we performed immunoprecipitation assays from cell lysates using anti Top2a antibodies. Western blot analysis of pull-downs confirms the direct interaction between Top2a and hPol I shown via its subunit RPA49. Furthermore, the observed signals for UBF are in line with an interaction in cells (Fig. 5B).

To clarify whether UBF-Top2a interaction is direct, we tested the binding of recombinant FLAG-tagged UBF (fUBF) and Top2a. Incubation of both proteins *in vitro* followed by a pulldown using anti-FLAG antibodies shows a clear band for Top2a in western blots (Fig. 5C, lane 2). Increasing salt concentration weakened (lane 3, 100 mM KCl) and finally abolished (lane 4, 200 mM KCl) the co-IP. We conclude that Top2a can interact with both, Pol I and UBF in human cells and *in vitro*.

## Discussion

The cryo-EM reconstruction of human Pol I demonstrates the overall conserved architecture of multi-subunit, DNA-dependent RNA polymerases in eukaryotes and completes the archive of yeast^17,18,20,91^ and mammalian^3–6^ nuclear Pol structures (Sup. Fig. 8). We find that human Pol I, like that of most organisms, carries a single-subunit stalk and built-in transcription factors show structural and functional similarities to TFIIF, TFIIE and TFIIS. Mapping of known hPol I mutations associated with human disease to the structural model (Sup. Fig. 6) rationalizes their effects on the enzyme.

We show that functional cross-species complementation of RPA49/34 subcomplexes is possible, which is in line with a conserved role in supporting initiation and elongation stages of the transcription cycle while accumulating divergent regulatory properties^92–94^. An increased flexibility of the clamp/stalk module in hPol I is indicated by the cryo-EM reconstruction (Fig. 2) and may explain an increased rate of incorrect nucleotide addition we observe in comparison to the yeast enzymes *in vitro* (Fig. 2B). This can be explained either by an impaired proof-reading due to reduced backtracking ability of hPol I, or a generally higher rate of substrate promiscuity. In yeast Pol I, module contraction is a feature of activation^95^. Especially during DNA melting upon transcription initiation^96,97^, contraction is required to stably associate melted template and non-template strands. Notably, the catalytic center, including the active site magnesium ion, is among the flexible parts in the hPol I cryo-EM reconstruction. The pronounced shelf module flexibility may indicate the importance of such a mechanism in higher eukaryotes, or simply point to a lack of defined intermediate conformations under close-to-native conditions in human cells.

While we do not observe any cryo-EM density for bound human RRN3, it can be assumed that binding to hPol I is similar to the *S. cerevisiae* counterpart^27–29^, due to sequence conservation of the factor^24^ and its binding sites in Pol I subunit RPA43 and the dock domain of subunit RPA1 (Sup. Data 1). Yeast Pol I subunit A14 is not involved in Rrn3 contacts^27–29^. Therefore, its absence in the human enzyme does not disagree with this model. Notably, purification by ion exchange chromatography leads to a dissociation of human RRN3 and the RPA49/34 heterodimer (Sup. Fig. 1B), indicating a reduced affinity and hence the possibility for efficient regulation of interaction with the core enzyme by PTMs, such as RRN3 phosphorylation^98^ and RPA49 acetylation^32^.

Most strikingly, our study identifies a previously unknown built-in transcription factor-like domain that resembles the fold of a truncated HMG box (Fig. 4). This ‘dock II’ domain is only found in higher organisms (Fig. 3) and shows similarities to HMG box 5 in UBF. While its function will be studied in more detail in the future, we find evidence that it may serve as an interaction platform for human Topoisomerase 2a. Three possible reasons for this interaction come to mind (Fig. 6): (1) Top2a could be part of Pol I initiation complexes in human cells^85^, while it does not appear to be involved in yeast PIC formation. Top2a recruitment to the downstream edge of human Pol I PICs via the built-in HMG box may be an attractive way to release tension from the DNA that accumulates upon spontaneous melting. In Pol II initiation systems, the XPB translocase in TFIIH occupies a similar position and carries out a comparable though not identical function in yeast^99^ and human PICs^100,101^. Deletion of the Top2a C-terminus leads to a six-fold reduction in RRN3 co-purification, but only a two-fold reduction in hPol I co-purification^85^, arguing for the possibility of RRN3 co-dependent Top2a recruitment via the foot-HMG box domain. (2) Positive supercoiling accumulates in the direction of transcription^102^, especially in Pol I-transcribed rDNA genes^103^, due to an increased loading rate^104^ and speed compared to other polymerases^39^. To release this supercoiling, Top2a may be recruited to the downstream face of elongating hPol I via the built-in HMG box. This may be reflected in an elongation-factor like behavior of Top2a and could be exclusive to the first round of transcription of a previously inactive rDNA gene. Following Top2a-supported opening of the gene by initial hPol I transcription, including nucleosome removal assisted by FACT^105^, association of UBF over the gene body^46^ may prevent closing and strong accumulation of positive supercoiling during subsequent rounds of Pol I transcription. (3) Co-dependent association of UBF with Top2a over the rDNA gene may create periodic hubs that allow the transient recruitment of Top2a to hPol I on active genes to release positive supercoiling. The three C-terminal HMG-boxes of UBF may be responsible for a hand-over of Top2a to the dock II-HMG box and recover Top2a following its transient interaction with Pol I. In addition, UBF association with DNA introduces additional supercoiling itself^106^. In actively transcribed genes, high on/off rates of UBF can be expected, leading to the local requirement of Top2a that could be satisfied by UBF association of the enzyme.

**Figure 6.**
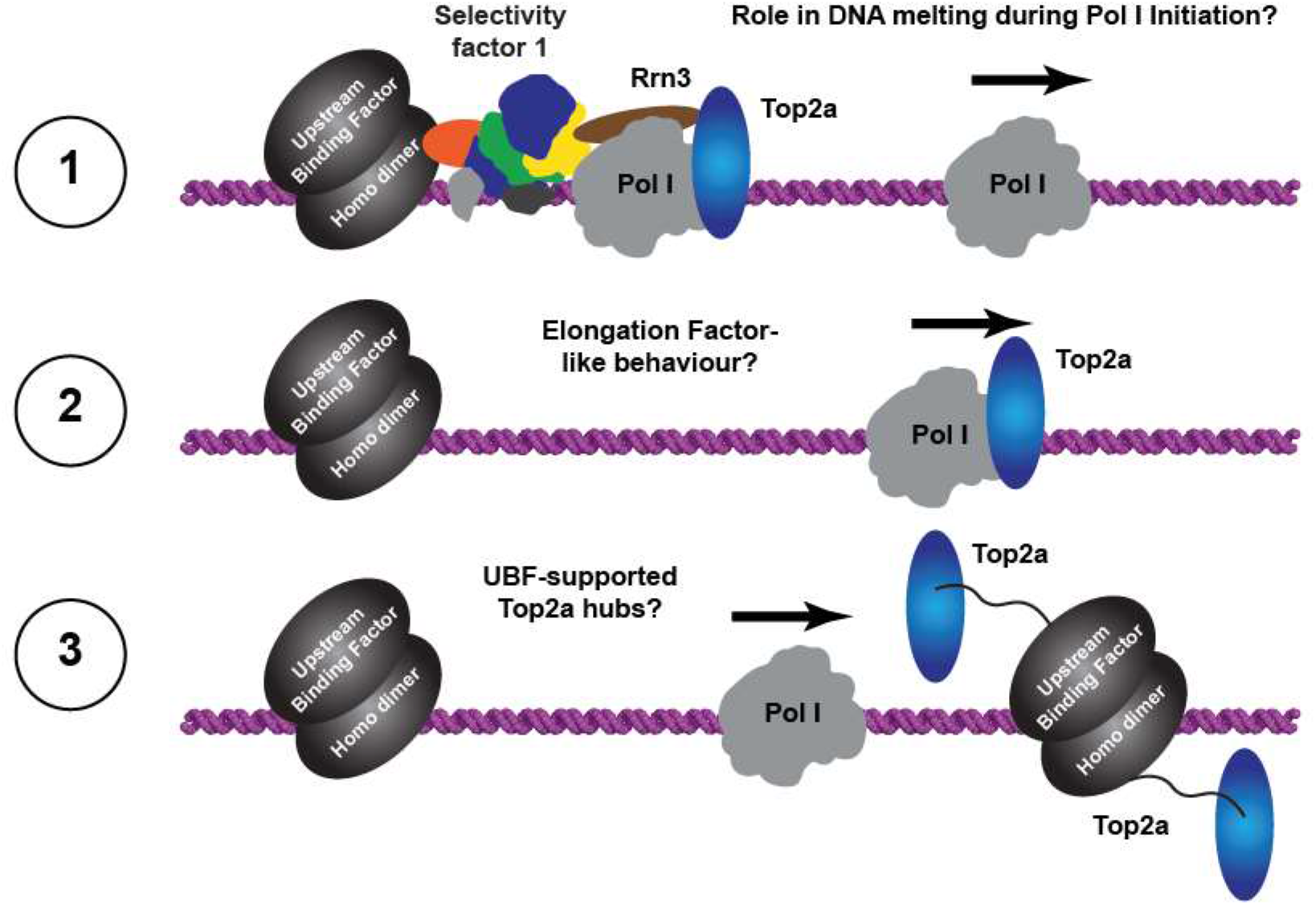
Possible roles of Top2a in human Pol I transcription. Three hypotheses are likely scenarios: (1) Top2a may support initiation by resolving supercoils generated during dsDNA melting. (2) Top2a may travel with Pol I in an elongation factor like manner to resolve positive supercoils upon their accumulation. (3) Supported by direct and indirect evidence, we speculate that UBF and Top2a cooperate to from ‘torsion release hubs’ at the 3’ region of the rDNA gene.

Options (2) and (3) are supported by fact that Top2a signal is detected on the entire gene and co-localization of Top2a with UBF in some regions is observed in ChIP-Seq studies (Fig. 5A). An initiation factor-like profile for Top2a that would point towards option (1) is not detected. Though possibly coincidental, further evidence for hypothesis (3) arises from phylogenetic analysis demonstrating that UBF versions start to appear in the same organism in which we detect the presence of the dock II domain (Sup. Fig. 9) and from the recent finding that Top2 localization to the nucleolus depends on Pol I activity in human cells^107^. In line with this, we demonstrate that physical interaction between UBF and Top2a is possible.

Nevertheless, additional functions for the HMG box-containing dock II domain independent of Top2a can be imagined. The domain clashes with the ‘trestle’ helix of the CTR9 subunit in the PAF-complex, a Pol II elongation factor^108^. This may prevent PAF action in human Pol I transcription, even though an effect in yeast Pol I elongation was reported^109^. Furthermore, the HMG-box containing SOX factors assist DNA-detachment from nucleosomes^110^ and SSRP1 is a component of FACT, that also contains a single HMG box and is required for hPol I transcription through nucleosomes^105^. In fact, single HMG box-containing proteins were described to functionally support human FACT^111^. Together with the positioning of dock II close to the incoming (downstream) DNA duplex, this also supports the speculation of a function in efficient nucleosome encounter of hPol I. Most of these factors, however, require a direct DNA-interaction of their HMG box, which appears unlikely for dock II due to occlusion of the DNA-interface by RPABC1 and its mutated DNA-binding site (Fig. 4).

During initial peer-review of this work, two groups also reported cryo-EM reconstructions of hPol I ^112,113^. The focus of one study lies on the structural basis of backtracking and cleavage^112^, while the other also reports a co-structure with RRN3^113^. Our colleagues present reconstructions with higher overall resolution, but do not comment on the role of the novel dock II domain and involvement of Top2a in rDNA transcription. Therefore, the findings of the three studies support and supplement each other.

While a detailed analysis of dock II function(s) will now commence, it is not surprising that another transcription factor-related domain is built into metazoan Pol I. In addition to TFIIF and TFIIE elements within subunits RPA49/34, TFIIS elements in RPA12 and a DNA-mimicking element in RPA1, integration of an HMG-box element seems to contribute to the accumulating specialization of Pol I during evolution.

## Author Contributions

JLD planned, carried out and evaluated all experiments including cell line generation, hPol I purification, functional biochemistry, structure determination and microscopy. MP collected cryo-EM data and contributed to functional biochemistry and cryo-EM processing. KS carried out phylogenetic analyses. ABl performed and evaluated fluorescence and confocal microscopy. MH and FBH purified proteins and contributed to functional biochemistry. ER, GA-P and AV contributed to data evaluation. VL supplied recombinant human Top2a. KP purified fUBF and carried out UBF-Top2 co-IPs. JLD, KT, CB and MP prepared and screened cryo-EM grids. ABr carried out mass spectrometry analysis. JCM and TM evaluated ChIP data. CE designed and supervised research and wrote the manuscript with input from all authors.

## Acknowledgements

The authors especially thank Philip Gunkel for his contribution. We thank all past and present members of the Engel lab, Achim Griesenbeck, Colyn Crane-Robinson, Christophe Lotz, Marlene Vayssieres, Klaus Grasser, Herbert Tschochner, and Philipp Milkereit for help and discussion, Gerhard Lehmann and Nobert Eichner for IT support, Joost Zomerdijk for UBF-constructs, Volker Cordes for the Hela P2 cell line, Remco Sprangers for shared cell culture, Dina Grohmann and the Archaea Center for fermentation, and Thomas Dresselhaus for access to fluorescence microscopes. This work was in part supported by the Emmy-Noether Programm (DFG grant no. EN 1204/1-1 to CE) of the German Research Council and Collaborative Research Center 960 (TP-A8 to CE).

## Data Availability

The cryo-EM density of human Pol I was deposited in the Electron Microscopy Data Bank. Model coordinates were deposited with the Protein Data Bank. Further material can be obtained from the corresponding author upon reasonable request.

## Methods

### CRISPR/Cas9 genome editing

HeLa cells were cultivated in DMEM medium (21885, Gibco) supplemented with 10 % FBS (10270, Gibco) and 1 % Penicillin/Streptomycin (P0781, Sigma Aldrich) at 37°C and 5 % CO_2_ atmosphere. Genomic integration of sfGFP ORF at the C-terminus of RPA1 was done by CRISPR/Cas9 according to a published protocol (Ran et al, 2013b) with some modifications and identical as previously published for RPAC1-sfGFP ^4^.

Design of the guide RNAs (gRNAs) was done with a web-based tool (https://www.benchling.com/crispr/) and annealed oligonucleotides (gRNA1 = GCTCCAAGGACCCTTGGTGA; gRNA2 = CGGGGTAGCTGCTATCTCAG) were cloned via BbsI as described in the manual into the Cas9n expression vector pSpCas9n(BB)-2A-Puro (PX462) V2.0, which was a gift from Feng Zhang (Addgene plasmid #62987; https://www.addgene.org/62987/; RRID: Addgene_62987). A donor plasmid carried a short GS-linker sequence with an embedded HRV 3C protease cleavage site and the sfGFP ORF surrounded by two large sequence segments homologous to the insertion locus in the genome.

HeLa cells were transfected with a 1:1:1 molar ratio of gRNA1 and gRNA2 vectors together with the donor plasmid using FuGENE HD Transfection Reagent (E2311, Promega) according to the manufacturer’s instructions. Several days later the GFP-expressing cells were enriched by flow cytometry using a BD FACSAria™ IIu cell sorter at the Central FACS Facility of the RCI Regensburg (Center for Interventional Immunology). GFP-positive cells were seeded as single cells on 96-well plates. After 2-3 weeks, colonies were expanded. These monoclonal populations were validated for the tag insertion by PCR on extracted genomic DNA (gDNA), sequencing and western blot.

About 1*10^6^ cell were resuspended in proteinase K buffer (20 mM Tris pH 7.5, 300 mM NaCl, 25 mM EDTA, 2 % (w/v) SDS, 0.2 ^mg^/_ml_ proteinase K) and incubated overnight at 50°C before performing isopropanol precipitation. The resuspended gDNA was used as template for PCR to validate the homozygous introduction of the GS-linker and sfGFP ORF into the POLR1A genomic locus (Primer: POLR1A-fwd1: 5’-TTGGGATCCGGTCAAACTC-3’, POLR1A-rev1: 5’-#CAGCAAAGCATGGCTTCC-3’, POLR1A-fwd2: 5’-CAGTGGGATCTTGGGATCTG-3’, POLR1A-rev2: 5’-TGCTACGCTGTACTTGACTC-3’). To further validate the result, the PCR product was gel extracted (QIAquick Gel Extraction Kit, 28706, QIAGEN) and sequenced (Microsynth Seqlab). Additional characterization of the selected homozygous cell line was done by Western Blot. Cells from a confluent 6 cm plate (about 2.7*10^6^ cells, 83.3901.300, Sarstedt) were harvested with 300 µl of boiling 1x SDS loading dye (3 % (w/v) glycerol, 1.68 % (v/v) β-mercaptoethanol, 0.03 % (w/v) bromophenol blue, 26 mM Tris pH 6.8, 0.42 % (w/v) SDS) and vigorously shaken at 95°C for 15 min. Prestained marker (7719S, NEB), as well as 10 µl of sample from the parental and the newly generated cell line, were loaded on an SDS gel (NP0223BOX, Thermo Fisher Scientific) and proteins were separated by electrophoresis. After blotting (Trans-Turbo Blot, Bio-Rad) the proteins onto a PVDF membrane (1704275, Bio-Rad), Ponceau S staining confirmed equal loading. The tagged protein RPA1 was detected by the primary antibody (sc-48385, Santa Cruz Biotechnology), which was subsequently detected by the fluorescently labeled secondary antibody (926-32210, Li-COR). Prestained marker and secondary antibody were detected by different wavelengths (Odyssey Infrared Imager Model 9120, Li-COR).

The selected cell line RPA1-sfGFP was cultivated adherently and adapted to suspension growth as follows: Cells from 8 flasks (about 7×10^7^ cells total; 83.3912.302, Sarstedt) were detached by incubation with trypsin (25300, Gibco) at 37°C for 5 min, transferred to a spinner flask (250 mL total volume; 4500, Corning) and cultured in suspension with high-glucose DMEM (11965, Gibco) supplemented with 1 % FBS (10270, Gibco) and 1 % Penicillin/Streptomycin (P0781, Sigma Aldrich) under moderate stirring at 37°C and 5 % CO_2_ atmosphere. To expand the culture, 1x the current volume of fresh media including all supplements was added when cells reached a density of ∼7×10^5 cells^/_ml_ and the culture was transferred to spinner flasks of increasing volume when required. Cells were harvested by centrifugation and washed with PBS before flash-freezing the pellet.

### Purification of human Pol I

Human Pol I purification was performed similarly to ^4^ with some modifications. RPA1-sfGFP cell pellet was resuspended in twice the volume of the cell pellet’s weight of lysis buffer (20 mM Hepes pH 7.8, 420 mM NaCl, 1 mM MgCl_2_, 10 µM ZnCl_2_, 0.5 % (v/v) NP-40, 4 mM β-mercaptoethanol, 1x protease inhibitor mix (Benzamidine & PMSF)) supplemented with 7 ^U^/_ml_ DNase I (M610A, Promega) and lysed by Dounce homogenization and incubation on ice for 30 min. After centrifugation at 20,000*g and 4°C for 15 min, the whole-cell lysate was incubated with pre-equilibrated GFP-Trap Dynabeads (gtd, Chromotek) for binding. The beads were washed once with four times and once with twice the slurry volume of wash buffer (20 mM Hepes pH 7.8, 420 mM NaCl, 1 mM MgCl_2_, 10 µM ZnCl_2_, 2 % (v/v) glycerol, 4 mM β-mercaptoethanol), before being eluted with the volume of the slurry with wash buffer supplemented with 10 µg of 3C protease per 1 g of cell pellet for 4 h at 4°C. In case an anion-exchange chromatography was performed, the GFP-elution was diluted with buffer A (20 mM Hepes pH 7.8, 1 mM MgCl_2_, 10 µM ZnCl_2_, 2 % (v/v) glycerol, 5 mM DTT) to reach a final concentration of 140 mM NaCl. The sample was loaded on a MonoQ 1.6/5 PC column (Pharmacia Biotech) with 60 mM ammonium sulfate and eluted stepwise in buffer A with increasing the concentration of ammonium sulfate up to 1 M. A linear gradient over five column volumes to 200 mM followed by steps of five column volumes with 200 mM, 350 mM, 600 mM and 1 M ammonium sulfate was applied. hPol I eluted at 350 mM ammonium sulfate concentration. hPol I was used immediately or flash-frozen in liquid nitrogen and stored at - 80°C for further experiments.

### RNA Elongation and Cleavage Assay

RNA Elongation and Cleavage Assay was performed as described ^4^ with small modifications. 0.5 pmol of Pol I from *S. cerevisiae, S. pombe*, or *H. sapiens* were preincubated with 0.25 pmol of different pre-annealed minimal or bubble nucleic acid scaffolds (sequence information summarized in Sup. Table 2 and schematically shown in each figure along with the gel)in transcription buffer (20 mM Hepes pH 7.8, 40 mM (NH_4_)_2_SO_4_, 28 mM NaCl, 8 mM MgSO_4_, 10 µM ZnCl_2_, 10 % (v/v) glycerol, 10 mM DTT) for 1 h at 20°C in a 45 µl reaction. In case purified RPA49/RPA34 heterodimer was added, 1x, 5x or 10x molar excess of heterodimer compared to polymerase was included during the preincubation. For RNA elongation, 10 µmol of each desired NTP (marked specifically at each lane in the figure) were added and the reaction was incubated for 1 h at 28°C. To examine cleavage activity, the preincubated reaction was incubated for 1 h at 28°C without the addition of NTPs. Afterwards, nucleic acid purification was examined by adding 5M NaCl to a final concentration of 0.5 M and 800 µl 100 % ethanol. After precipitation for at least 1 h at -20°C, the sample was centrifuged for 30 min at 20,000*g and 4°C. The pellet was washed with 80 % ethanol and, after drying, resuspended in 1x RNA loading dye (4 M Urea, 1x TBE, 0.01% bromophenol blue and 0.01% xylene cyanol only for FAM-labeled constructs). The sample was heated to 95°C for 5 min. As control 0.25 pmol of scaffold were treated identically, without addition of polymerase and NTPs. 0.125 pmol of FAM-labeled RNA product were separated by gel electrophoresis (20 % polyacrylamide gel containing 7 M Urea) and visualized with a Typhoon FLA9500 (GE Healthcare).

### Purification of RPA49/RPA34 variants

The *S. cerevisiae* full-length heterodimer was purified as described^19^. Sc A49 with a C-terminal hexa-histidine tag and Sc A34 were co-expressed in E. coli BL21 (DE3) RIL in LB medium with 0.2 mM IPTG for 18 h at 18°C. The cells were resupended in lysis buffer (50 mM Tris pH 7.5, 300 mM NaCl, 10 mM β-mercaptoethanol, 1x protease inhibitor (PI) mix (Benzamidine & PMSF)) and sonified. After centrifugation, the lysate was loaded onto preequilibrated Ni-NTA beads (30230, Qiagen) by gravity-flow, washed with six times the bed volume of buffer Wash I (50 mM Tris pH 7.5, 1 M NaCl, 10 mM β-mercaptoethanol, 1x PI) and six times the bed volume of Wash II (50 mM Tris pH 7.5, 300 mM NaCl, 30 mM imidazole, 10 mM β- mercaptoethanol, 1x PI) before elution (50 mM Tris pH 7.5, 300 mM NaCl, 100 mM imidazole, 10 mM β-mercaptoethanol, 1x PI). The sample was diluted 3-fold with dilution buffer (50 mM Tris pH 7.5, 10 mM β-mercaptoethanol) before loading onto a MonoS 5/50 GL column (GE Healthcare) with buffer A (50 mM Tris pH 7.5, 100 mM NaCl, 5 mM DTT). Elution was performed with a linear gradient of NaCl concentration up to 1 M. Sc A49/34 eluted at around 280 mM NaCl. The corresponding fractions were pooled and concentrated with 10 kDa cut off (UFC801024, Millipore) and applied to a Superdex200 Increase 100/300 (GE Healthcare) equilibrated with buffer A. Pooled peak fractions were concentrated and flash-frozen for storage at -80°C.

The different variants of the human heterodimer (RPA49^FL^/RPA34^FL^, RPA49^FL^/RPA34^1-343^, RPA34^131-510^) were cloned with a N-terminal 6xHis-tag on RPA49 and untagged RPA34, except for RPA34^131-510^, which carries an N-terminal 6xHis-tag itself. The proteins were coexpressed in *E. coli* BL21 (DE3) RIL in LB medium with 0.2 mM IPTG overnight at 18°C. Cells were resuspended in lysis buffer (50 mM MES pH 6.3, 300 mM NaCl, 10 mM β-mercaptoethanol, 1x protease inhibitor (PI) mix (Benzamidine & PMSF)) and lysed by sonification. After centrifugation, the lysate was loaded onto preequilibrated Ni-NTA beads (30230, Qiagen) by gravity-flow, washed subsequently with six times the bed volume of buffer Wash I (50 mM MES pH 6.3, 1 M NaCl, 10 mM β-mercaptoethanol, 1x PI), ATP-Wash (50 mM MES pH 6.3, 1 M NaCl, 10 mM β-mercaptoethanol, 1x PI supplemented with 2 ^mg^/_ml_ denatured proteins and 0.5 mM ATP), another ATP-Wash after 10 min of incubation and Wash II (50 mM MES pH 6.3, 300 mM NaCl, 10 mM imidazole, 10 mM β-mercaptoethanol, 1x PI) before elution (50 mM MES pH 6.3, 300 mM NaCl, 200 mM imidazole, 10 mM β-mercaptoethanol, 1x PI). The ATP-Wash steps were performed at room temperature. The sample was diluted 5-fold with buffer A (50 mM Tris pH 7.5, 10 mM β-mercaptoethanol) before loading onto a MonoS 5/50 GL column (GE Healthcare) with buffer A supplemented with 100 mM NaCl. Elution was performed with a linear gradient of NaCl concentration up to 2 M. The corresponding fractions were pooled and concentrated with 10kDa cut off (UFC801024, Millipore) and applied to a Superdex200 Increase 100/300 (GE Healthcare) equilibrated with SEC buffer (50 mM Tris pH 7.5, 150 mM NaCl, 5 mM DTT). Pooled peak fractions were concentrated and flash-frozen for storage at -80°C.

### Purification of recombinant dock II domain

Two variants of the human dock II domain (RPA1^1060-1155^ (full-length), RPA1^1081-1146^ (minimal)) were cloned with a C-terminal His-MBP-tag. The proteins as well as tag-only were expressed overnight at 20°C in E. coli BL21 (DE3) RIL in LB medium with 0.2 mM IPTG. Cells were resuspended in lysis buffer (50 mM MES pH 6.3, 300 mM NaCl, 10 mM β-mercaptoethanol, 1x protease inhibitor (PI) mix (Benzamidine & PMSF)) and lysed by sonification. After centrifugation, the lysate was loaded onto preequilibrated Ni-NTA beads (30230, Qiagen) by gravity-flow, washed subsequently with six times the bed volume of buffer Wash I (50 mM MES pH 6.3, 1 M NaCl, 10 mM β-mercaptoethanol, 1x PI) and Wash II (50 mM MES pH 6.3, 300 mM NaCl, 10 mM imidazole, 10 mM β-mercaptoethanol, 1x PI) before elution (50 mM MES pH 6.3, 300 mM NaCl, 200 mM imidazole, 10 mM β-mercaptoethanol, 1x PI). The eluat was buffer-exchanged to SEC buffer (50 mM Tris pH 7.5, 150 mM NaCl, 5 mM DTT) with a PD10 column (17-0850-01, GE Healthcare) and applied to a Superdex 75 Increase 10/300 GL(GE Healthcare) equilibrated with SEC buffer. Pooled peak fractions were concentrated and flash-frozen for storage at -80°C.

### Electrophoretic mobility shift assay

A total of 100 fmol pre-annealed 40 bp DNA (EMSA-DNA-strand1: 5’-Cy5-CTGGAACAACACTCAACCCTATCTCGGTCTATTCTTTTGA-3’; EMSA-DNA-strand2: 5’-TCAAAAGAATAGACCGAGATAGGGTTGAGTGTTGTTCCAG -3’) were mixed with up to 50-fold molar excess of purified protein (as labeled in the figure) in EMSA buffer 1 or 2 (EMSA-buffer-1: 10 mM Tris pH 7.5, 50 mM NaCl, 1 mM MgCl_2_, 4 % glycerol, 0.5 mM EDTA, 0.5 mM DTT; EMSA-buffer-2: 20 mM Hepes pH 7.8, 150 mM NaCl, 2 % glycerol, 0. 2% Triton-100, 0.2 % Tween-20, 5 mM DTT) and incubated at room temperature for 30 min. Afterwards 6x loading dye (10 mM Tris pH 7.6, 60 mM EDTA, 60 % glycerol, 0.03 % Orange G) was added to reach 1x concentration. 10% polyacrylamide gels in 0.4x TBE were pre-run at 110 V for 30 min before the reaction was separated at 110 V for 1:45 h at 4°C. The Cy5-labeled DNA was detected with a Typhoon FLA9500 (GE Healthcare).

### Confocal microscopy

For fluorescence imaging, cells were grown adherently on glass cover slips to 50 % confluency. After washing the cells with pre-warmed (37°C) PBS, they were fixed with 3.7 % paraformaldehyde in PBS for 10 min at 37°C. The fixation was stopped by replacing the solution with 100 mM glycine in PBS for 5 min at 37°C. After that cells were washed twice with PBS, mounted on the specimen slide with the help of a drop of Prolong Gold Antifade Mountant with DAPI (P36941, Thermo Fisher Scientific), and dried in the dark at least overnight.

The fluorescent specimens were imaged using a Plan-Apochromat 63x/1,4 Oil DIC Objective at a Zeiss LSM980/Airyscan 2 confocal microscope. sfGFP was excited by a 488 nm diode laser and emission was detected using a 300-720 nm band pass filter. Separately DAPI was excited by a 405 nm diode laser and emission was detected using a 300-720 nm band pass filter. For the 3D model a Z-stack was imaged using the internal GaAsP-PMT detectors from 490-668 nm for sfGFP and 410-473 nm for DAPI in a two-track process. Image processing was done using the Zeiss AxioVision software. The 3D Volume images were created in Imaris 9.6.

### Analysis of Pol I subunits RPA1, RPA34, RPA43 and A14

Data sets from Pol I subunits were generated using their corresponding InterPro^114^ entries (RPA1: IPR015699, RPA34: IPR013240, RPA43: IPR041901 and IPR041178, A14: IPR013239 downloaded on 07.06.2021). A common dataset of RPA1, RPA34 and RPA43 was generated by searching for common species within the three InterPro families. To each obtained species the concatenated sequence of RPA1, RPA34 and RPA43 was assigned.

### Phylogenetic analysis

Sequence alignment tool MAFFT ^115^ has been used with default options and a gap open penalty of 70. The resulting alignment was filtered manually on highly diverged sequences. To improve the quality of the phylogenetic analysis without losing information for each genus only one sequence was chosen. On the resulting data set with 513 sequences Gblocks ^116^ (options: b3=5000, b4=2, b5=a) has been applied to remove uninformative columns. By means of RAxML ^117^ using the option -f a and the substitution model PROTGAMMAAUTO 100 trees were generated and a consensus tree was derived. The root has been placed between the supergroups of Sar and Haptophyta and the supergroup of Amorphea ^118^. The resulting phylogenetic tree was analyzed with respect to the taxonomic distribution. Sequences were grouped according to branching points in the phylogenetic tree (Fig. 3). In order to retrieve the taxonomic group where the A14 subunit is present the species related to the A14 subunit InterPro entry are compared with the species given in the phylogenetic tree.

### Sequence analysis of RPA34 and RPA1

By means of MAFFT sequence alignment of each subunit was generated using varied gap open penalties (RPA34: 50, RPA1: 20). Due to higher sequence variety within RPA34 sequences BLOSUM30 was used instead of the default parameter. In order to account the divergence between the taxonomic groups given from the phylogenetic tree, the alignment was split into these groups and each group was analyzed separately on the presence or absence of the RPA34 C-terminal extension, the RPA1 foot domain and the RPA1 expander domain. Sequences from *Homo sapiens* have been used as reference to identify the region of interests (399-510; 1074-1139; 1365-1488, respectively). The median length and standard deviation of the regions of interest have been calculated for each group. To unravel the sequence and structural conservation of the regions of interest the conservation score given in Jalview ^119^ has been extracted after removing all columns containing only gaps. The mean conservation score is calculated by summing up over all column scores divided by the number of columns. Scores are grouped into 5 categories: not conserved (0-3), weakly conserved (3-5), medium conserved (5-7), conserved (7-9), strongly conserved (9-11). Secondary structures were predicted using Ali2D ^79,120^. Secondary structure elements were assigned when more than 5 amino acids have medium to high probability in more than 90 % of the sequences within each group. Bridging of two secondary structure elements over less than 5 differently annotated amino acids are counted as one element. If gaps are present in more than 90 % of the sequences, they are ignored.

### Mass Spectrometry

Protein bands were cut out from the gel, washed with 50 mM NH_4_HCO_3_, 50 mM NH_4_HCO_3_/acetonitrile (3/1), 50 mM NH_4_HCO_3_/acetonitrile (1/1) and lyophilized. After a reduction/alkylation treatment and additional washing steps, proteins were *in gel* digested with trypsin (Trypsin Gold, mass spectrometry grade, Promega) overnight at 37 °C. The resulting peptides were sequentially extracted with 50 mM NH_4_HCO_3_ and 50 mM NH_4_HCO_3_ in 50 % acetonitrile. After lyophilization, peptides were reconstituted in 20 µl 1 % TFA and separated by reversed-phase chromatography. An UltiMate 3000 RSLCnano System (Thermo Fisher Scientific, Dreieich) equipped with a C18 Acclaim Pepmap100 preconcentration column (100 µm i.D. x 20 mm, Thermo Fisher Scientific) and an Acclaim Pepmap100 C18 nano column (75 µm i.d. x 250 mm, Thermo Fisher Scientific) was operated at a flow rate of 300 ^nl^/_min_ and a 60 min linear gradient of 4 % to 40 % acetonitrile in 0.1 % formic acid. The LC was online-coupled to a maXis plus UHR-QTOF System (Bruker Daltonics) via a CaptiveSpray nanoflow electrospray source. Acquisition of MS/MS spectra after CID fragmentation was performed in data-dependent mode at a resolution of 60,000. The precursor scan rate was 2 Hz processing a mass range between ^m^/_z_ 175 and ^m^/_z_ 2000. A dynamic method with a fixed cycle time of 3 s was applied via the Compass 1.7 acquisition and processing software (Bruker Daltonics). Prior to database searching with Protein Scape 3.1.3 (Bruker Daltonics) connected to Mascot 2.5.1 (Matrix Science), raw data were processed in Data Analysis 4.2 (Bruker Daltonics). Swiss-Prot *Homo sapiens* database (release-2020_01, 220420 entries) was used for database search with the following parameters: enzyme specificity trypsin with one missed cleavage allowed, precursor tolerance 0.02 Da, MS/MS tolerance 0.04 Da, Mascot peptide ion-score cut-off 25. Deamidation of asparagine and glutamine, oxidation of methionine, carbamidomethylation or propionamide modification of cysteine were set as variable modifications.

### Native PAGE

To investigate protein-protein interaction, blue-native PAGE was performed. Five times molar excess of MBP only or tagged human dock II domain was incubated with recombinant Top2a ΔC (1-1217) in binding buffer (20 mM Hepes pH 8.0, 150 mM NaCl, 50 mM KCl, 1 mM MgCl2, 2 % glycerol, 2 mM β-mercaptoethanol) for 30 min at room temperature. After adding NativePAGE sample buffer, the samples were separated on a Native PAGE 3-12% gradient gel at 150 V for 90 min with light blue cathode and anode buffer (NativePAGE™ Novex® Bis-Tris Gel System, BN1003BOX, Novex) and coomassie stained.

### Top2a co-immunoprecipitation

To investigate Top2a interaction partners, co-immunoprecipitation was performed from U2OS Nuclear Extract (15 mg/ml total protein). Top2a was immuno-precipitated using an anti-Top2a antibody (ab12318, Abcam) immobilized on Dynabeads Protein A magnetic beads (Thermofisher, c/n 10001D) according to the manufacturer’s instruction. Antibodies were cross-linked to beads using DPM (Thermofisher, c/n 21666) as recommended by the manufacturer. Beads were blocked with BSA in PBS overnight. 100 µL NE was diluted by dilution buffer (25 mM TrisHCl pH7.9, 12.5 mM MgCl_2_, 10% glycerol, 0.03% NP40) to a final KCl concentration of 150 mM and treated by 500 U of benzonase (Sigma, E1014) for 30 min at 4 ^°^C. 25 µl of the beads were added and the suspension was incubated on a rotating wheel for 1 hour at 4°C. Beads were washed three times with 100 µl wash buffer (25 mM TrisHCl pH7.9, 150 mM KCl, 12.5 mM MgCl_2_, 10% glycerol, 0.03% NP40) and proteins were eluted by incubation in 1x LDS-sample buffer (Thermofisher, c/n NP0007) at 65°C for 10 minutes. Immunoprecipitated proteins were analyzed by Western blot using anti-UBF, anti-RPA49 and anti-Top2a antibodies (sc-9131, Santa Cruz; 611413 BD Transduction; and ab12318, Abcam).

### UBF-Top2a pulldown

To investigate protein-protein interaction, a pulldown assay using purified recombinant Flag-tagged UBF (fUBF) and purified Top2a was performed. fUBF was expressed in insect cells and purified as described earlier^121^. Top2a was obtained from Inspiralis (c/n HT210). Proteins were incubated together in pulldown buffer (25 mM TrisHCl pH7.9, 12.5 mM MgCl_2_, 10% glycerol, 0.03% NP40 supplemented with 50,100 or 200 mM KCl as marked in the Fig. 5C) for 20 min at 4°C. To each sample, 20 µl anti-FLAG M2 Magnetic Beads (Sigma, M8823) were added and the suspension was incubated on a rotating wheel for 30 min at 4°C. Beads were washed three times with wash buffer (25 mM TrisHCl pH7.9, 12.5 mM MgCl_2_, 10% glycerol, 0.03% NP40 supplemented with 50,100 or 200 mM KCl) and proteins were eluted by incubation in 1x LDS-sample buffer (Thermofisher, c/n NP0007) at 65°C for 10 minutes. Proteins were analyzed by Western Blot using anti-UBF and anti-Top2a antibodies (sc-9131, Santa Cruz; ab12318, Abcam).

### Reanalysis of previously published ChIP datasets

Raw data was handled, mapping coordinates exacted, and the data displayed as previously published ^90^. The used data was: Top2A GSE99197_SRR5585950_TOP2A-MEF ^87^. ArrayExpress E-MTAB-5839 data sets were: ChIP-seq_UBF_MEFs_UBFfl_Rep1; ChIP-seq_RPI_MEFs_UBFfl_Rep1; ChIP-seq_Rrn3_MEFs_UBFfl_Rep1; ChIP-seq_TBP_MEFs_UBFfl_Rep1; ChIP-seq_TAF68_MEFs_UBFfl_Rep1 ^46^. Taf1c is not included in the Figure since it is identical to the Taf1b mapping but data is also available in E-MTAB-5839 as ChIP-seq_TAF95_MEFs_UBFfl_Rep1.

### Negative stain EM

hPol I samples were centrifuged (4°C; 15,000 rpm; Eppendorf table top centrifuge) for 5 min. Five µl of the samples were then applied to glow-discharged 400-mesh copper grids (G2400C; Plano) with a self-made carbon film of ∼7 nm thickness (Pilsl, et al, Methods Mol Biol, in press). After 1 min, grids were washed in ddH_2_O for 30 s, and stained three times with 5 µl saturated uranyl formiate solution (2x 20s, 1x 30 s). After each step, excess liquid was removed with a filter paper. Images were collected on a JEOL 2100-F Transmission Electron Microscope operated at 200 keV and equipped with TVIPS-F416 (4kx4k) CMOS-detector at 40,000x magnification (pixel size 2.7 Å) with alternating defocus (−1 to -3 µm).

The images were processed using RELION 3.1^57^ as shown in Sup. Fig 1. A total of 76 micrographs were analyzed, yielding 46,196 auto-picked particles using Laplacian-of-Gaussian (LoG) routine. Following reference-free 2D sorting, a 3D classification (reference PDB: 5M3M low-pass filtered to 60 Å) yielded three reconstructions with different clamp/stalk flexibilities (Sup. Fig. 1).

### Cryo-EM grid preparation and data collection

Reconstructions suffered from poor Fourier completeness. Screening for suitable conditions using crosslinking, gradient fixation^122^ and detergents, or variation of grid support types graphene (-oxide), ultrathin carbon or gold foil (Pilsl et al., Methods Mol Biol, in press) had limited success in removing orientational bias. Tilted data collection partially improved the bias even though 3D reconstruction was still hampered. Nevertheless, best results were obtained with GFP-trap eluted sample directly applied to graphene-oxide supported grids. However, this strategy retains some remaining 3C protease in the sample (Fig. 1C; Sup. Fig. 1A) that may have a negative influence on signal-to-noise ratio.

Graphene oxide grids were prepared using the surface assembly method on Quantifoil R1.2/1.3 grids^123^. Three microliters of sample were applied and incubated for 30 s at 100 % humidity at 4°C in a Vitrobot mark IV, blotted for 3 s with blot-force 8 and plunged into liquid ethane. A total of 9,709 micrograph movies were collected on a CryoArm200 cryo-electron microscope (JEOL) equipped with a K2 direct electron detector (Gatan), in-column energy filter and Cold-Field Emission Gun (low-flash interval 4 h). A total dose of 40 e^-^/A^2^ was fractionated over 40 frames at a defocus range of -1.2 to -2.7 µm using SerialEM^124^ in a 5×5 multi-hole strategy as described^60^.

### Cryo-EM image processing and model building

Pre-processing was carried out using WARP^56^, followed by 2D and 3D classification and auto-refinement using Relion 4.0^57^. During pre-processing motion-correction, CTF estimation and particle picking was performed. The pixel size was binned to 1.50846 Å/pix and particles extracted with a box size of 190. Rough 2D classification followed by 3D classification using a reference of hPol I obtained after stringent 2D classification and 3D refinement yielded a reconstruction at an overall resolution of 4.09 Å. Further 3D classification was performed to investigate the occupancy and flexibility of the dimerization domain of RPA49/34 and the clamp/stalk region. Models for common subunits RPABC1-5 and the RPAC1/2 assembly were transferred from a hPol III reconstruction^5^. Homology models of the hPol I subunits RPA1, RPA2, RPA49, RPA34, RPA12 and RPA43 were generated based on sequence and secondary structure alignments with the crystal structures of their *S. cerevisiae* counterparts (Sup. Data 1) using the MODELLER software package^58^. The models were adjusted in COOT^125^ and real-space refined using Phenix^126^. At later stages, released AlphaFold^59^ models were used to guide chain-tracing in poorly resolved areas. A model of the stalk subunit RPA43 is included in some figures, but was not deposited due to poor or absent cryo-EM density resulting from flexibility.

## References

1. Roeder, R. G. & Rutter, W. J. Multiple forms of DNA-dependent RNA polymerase in eukaryotic organisms. Nature 224, 234–237 (1969).

2. Vannini, A. & Cramer, P. Conservation between the RNA polymerase I, II, and III transcription initiation machineries. Molecular cell 45, 439–446; 10.1016/j.molcel.2012.01.023 (2012).

3. Bernecky, C., Herzog, F., Baumeister, W., Plitzko, J. M. & Cramer, P. Structure of transcribing mammalian RNA polymerase II. Nature 529, 551–554; 10.1038/nature16482 (2016).

4. Ramsay, E. P. et al. Structure of human RNA polymerase III. Nature communications 11, 6409; 10.1038/s41467-020-20262-5 (2020).

5. Girbig, M. et al. Cryo-EM structures of human RNA polymerase III in its unbound and transcribing states. Nature structural & molecular biology 28, 210–219; 10.1038/s41594-020-00555-5 (2021).

6. Wang, Q. et al. Structural insights into transcriptional regulation of human RNA polymerase III. Nature structural & molecular biology 28, 220–227; 10.1038/s41594-021-00557-x (2021).

7. Li, L. et al. Structure of human RNA polymerase III elongation complex. Cell research 31, 791–800; 10.1038/s41422-021-00472-2 (2021).

8. Goodfellow, S. J. & Zomerdijk, J. C. B. M. Basic mechanisms in RNA polymerase I transcription of the ribosomal RNA genes. Sub-cellular biochemistry 61, 211–236; 10.1007/978-94-007-4525-4_10 (2013).

9. Klinge, S. & Woolford, J. L. Ribosome assembly coming into focus. Nature reviews. Molecular cell biology 20, 116–131; 10.1038/s41580-018-0078-y (2019).

10. Warner, J. R. The economics of ribosome biosynthesis in yeast. Trends in biochemical sciences 24, 437–440 (1999).

11. Hannan, K. M., Sanij, E., Rothblum, L. I., Hannan, R. D. & Pearson, R. B. Dysregulation of RNA polymerase I transcription during disease. Biochimica et biophysica acta 1829, 342–360; 10.1016/j.bbagrm.2012.10.014 (2013).

12. Ferreira, R., Schneekloth, J. S., Panov, K. I., Hannan, K. M. & Hannan, R. D. Targeting the RNA Polymerase I Transcription for Cancer Therapy Comes of Age. Cells 9; 10.3390/cells9020266 (2020).

13. Sanij, E. et al. CX-5461 activates the DNA damage response and demonstrates therapeutic efficacy in high-grade serous ovarian cancer. Nature communications 11; 10.1038/s41467-020-16393-4 (2020).

14. Mars, J.-C. et al. The chemotherapeutic agent CX-5461 irreversibly blocks RNA polymerase I initiation and promoter release to cause nucleolar disruption, DNA damage and cell inviability. NAR cancer 2, zcaa032; 10.1093/narcan/zcaa032 (2020).

15. Jacobs, R. Q., Huffines, A. K., Laiho, M. & Schneider, D. A. The small molecule BMH-21 directly inhibits transcription elongation and DNA occupancy of RNA polymerase I in vivo and in vitro. The Journal of biological chemistry, 101450; 10.1016/j.jbc.2021.101450 (2021).

16. Russell, J. & Zomerdijk, J. C. B. M. The RNA polymerase I transcription machinery. Biochemical Society symposium, 203–216 (2006).

17. Engel, C., Sainsbury, S., Cheung, A. C., Kostrewa, D. & Cramer, P. RNA polymerase I structure and transcription regulation. Nature 502, 650–655; 10.1038/nature12712 (2013).

18. Fernández-Tornero, C. et al. Crystal structure of the 14-subunit RNA polymerase I. Nature 502, 644–649; 10.1038/nature12636 (2013).

19. Geiger, S. R. et al. RNA polymerase I contains a TFIIF-related DNA-binding subcomplex. Molecular cell 39, 583–594; 10.1016/j.molcel.2010.07.028 (2010).

20. Hoffmann, N. A. et al. Molecular structures of unbound and transcribing RNA polymerase III. Nature 528, 231–236; 10.1038/nature16143 (2015).

21. Grummt, I. Life on a planet of its own: regulation of RNA polymerase I transcription in the nucleolus. Genes & development 17, 1691–1702; 10.1101/gad.1098503R (2003).

22. Mayer, C., Zhao, J., Yuan, X. & Grummt, I. mTOR-dependent activation of the transcription factor TIF-IA links rRNA synthesis to nutrient availability. Genes & development 18, 423–434; 10.1101/gad.285504 (2004).

23. Zhao, J., Yuan, X., Frödin, M. & Grummt, I. ERK-dependent phosphorylation of the transcription initiation factor TIF-IA is required for RNA polymerase I transcription and cell growth. Molecular cell 11, 405–413 (2003).

24. Blattner, C. et al. Molecular basis of Rrn3-regulated RNA polymerase I initiation and cell growth. Genes & development 25, 2093–2105; 10.1101/gad.17363311 (2011).

25. Bodem, J. et al. TIF-IA, the factor mediating growth-dependent control of ribosomal RNA synthesis, is the mammalian homolog of yeast Rrn3p. EMBO reports 1, 171–175; 10.1038/sj.embor.embor605 (2000).

26. Moorefield, B., Greene, E. A. & Reeder, R. H. RNA polymerase I transcription factor Rrn3 is functionally conserved between yeast and human. Proceedings of the National Academy of Sciences of the United States of America 97, 4724–4729; 10.1073/pnas.080063997 (2000).

27. Engel, C., Plitzko, J. & Cramer, P. RNA polymerase I-Rrn3 complex at 4.8 Å resolution. Nature communications 7, 12129; 10.1038/ncomms12129 (2016).

28. Pilsl, M. et al. Structure of the initiation-competent RNA polymerase I and its implication for transcription. Nature communications 7, 12126; 10.1038/ncomms12126 (2016).

29. Torreira, E. et al. The dynamic assembly of distinct RNA polymerase I complexes modulates rDNA transcription. eLife 6; 10.7554/eLife.20832 (2017).

30. Peyroche, G. et al. The recruitment of RNA polymerase I on rDNA is mediated by the interaction of the A43 subunit with Rrn3. The EMBO journal 19, 5473–5482; 10.1093/emboj/19.20.5473 (2000).

31. Fath, S., Kobor, M. S., Philippi, A., Greenblatt, J. & Tschochner, H. Dephosphorylation of RNA polymerase I by Fcp1p is required for efficient rRNA synthesis. The Journal of biological chemistry 279, 25251–25259; 10.1074/jbc.M401867200 (2004).

32. Chen, S. et al. Repression of RNA polymerase I upon stress is caused by inhibition of RNA-dependent deacetylation of PAF53 by SIRT7. Molecular cell 52, 303–313; 10.1016/j.molcel.2013.10.010 (2013).

33. Comai, L., Tanese, N. & Tjian, R. The TATA-binding protein and associated factors are integral components of the RNA polymerase I transcription factor, SL1. Cell 68, 965–976 (1992).

34. Engel, C. et al. Structural Basis of RNA Polymerase I Transcription Initiation. Cell 169, 120-131.e22; 10.1016/j.cell.2017.03.003 (2017).

35. Merkl, P. E. et al. RNA polymerase I (Pol I) passage through nucleosomes depends on Pol I subunits binding its lobe structure. The Journal of biological chemistry 295, 4782–4795; 10.1074/jbc.RA119.011827 (2020).

36. Scull, C. E., Lucius, A. L. & Schneider, D. A. The N-terminal domain of the A12.2 subunit stimulates RNA polymerase I transcription elongation. Biophysical journal 120, 1883–1893; 10.1016/j.bpj.2021.03.007 (2021).

37. Kuhn, C.-D. et al. Functional architecture of RNA polymerase I. Cell 131, 1260–1272; 10.1016/j.cell.2007.10.051 (2007).

38. Turowski, T. W. et al. Nascent Transcript Folding Plays a Major Role in Determining RNA Polymerase Elongation Rates. Molecular cell 79, 488-503.e11; 10.1016/j.molcel.2020.06.002 (2020).

39. Lisica, A. et al. Mechanisms of backtrack recovery by RNA polymerases I and II. Proceedings of the National Academy of Sciences of the United States of America 113, 2946–2951; 10.1073/pnas.1517011113 (2016).

40. Merkl, P. et al. Binding of the termination factor Nsi1 to its cognate DNA site is sufficient to terminate RNA polymerase I transcription in vitro and to induce termination in vivo. Molecular and cellular biology 34, 3817–3827; 10.1128/MCB.00395-14 (2014).

41. Reiter, A. et al. The Reb1-homologue Ydr026c/Nsi1 is required for efficient RNA polymerase I termination in yeast. The EMBO journal 31, 3480–3493; 10.1038/emboj.2012.185 (2012).

42. Moss, T., Langlois, F., Gagnon-Kugler, T. & Stefanovsky, V. A housekeeper with power of attorney: the rRNA genes in ribosome biogenesis. Cellular and molecular life sciences : CMLS 64, 29–49; 10.1007/s00018-006-6278-1 (2007).

43. Gorski, J. J. et al. A novel TBP-associated factor of SL1 functions in RNA polymerase I transcription. The EMBO journal 26, 1560–1568; 10.1038/sj.emboj.7601601 (2007).

44. Denissov, S. et al. Identification of novel functional TBP-binding sites and general factor repertoires. The EMBO journal 26, 944–954; 10.1038/sj.emboj.7601550 (2007).

45. Kwon, H. & Green, M. R. The RNA polymerase I transcription factor, upstream binding factor, interacts directly with the TATA box-binding protein. The Journal of biological chemistry 269, 30140–30146 (1994).

46. Herdman, C. et al. A unique enhancer boundary complex on the mouse ribosomal RNA genes persists after loss of Rrn3 or UBF and the inactivation of RNA polymerase I transcription. PLoS genetics 13, e1006899; 10.1371/journal.pgen.1006899 (2017).

47. Gunkel, P., Iino, H., Krull, S. & Cordes, V. C. ZC3HC1 Is a Novel Inherent Component of the Nuclear Basket, Resident in a State of Reciprocal Dependence with TPR. Cells 10, 1937; 10.3390/cells10081937 (2021).

48. Hannan, R. D. et al. Affinity purification of mammalian RNA polymerase I. Identification of an associated kinase. The Journal of biological chemistry 273, 1257–1267; 10.1074/jbc.273.2.1257 (1998).

49. Jacobs, R. Q., Ingram, Z. M., Lucius, A. L. & Schneider, D. A. Defining the divergent enzymatic properties of RNA polymerases I and II. The Journal of biological chemistry 296, 100051; 10.1074/jbc.RA120.015904 (2021).

50. Engel, C., Neyer, S. & Cramer, P. Distinct Mechanisms of Transcription Initiation by RNA Polymerases I and II. Annual review of biophysics 47, 425–446; 10.1146/annurev-biophys-070317-033058 (2018).

51. Fernández-Tornero, C. RNA polymerase I activation and hibernation: unique mechanisms for unique genes. Transcription 9, 248–254; 10.1080/21541264.2017.1416267 (2018).

52. Kostrewa, D., Kuhn, C.-D., Engel, C. & Cramer, P. An alternative RNA polymerase I structure reveals a dimer hinge. Acta crystallographica. Section D, Biological crystallography 71, 1850–1855; 10.1107/S1399004715012651 (2015).

53. Neyer, S. et al. Structure of RNA polymerase I transcribing ribosomal DNA genes. Nature; 10.1038/nature20561 (2016).

54. Tafur, L. et al. Molecular Structures of Transcribing RNA Polymerase I. Molecular cell 64, 1135–1143; 10.1016/j.molcel.2016.11.013 (2016).

55. Tafur, L. et al. The cryo-EM structure of a 12-subunit variant of RNA polymerase I reveals dissociation of the A49-A34.5 heterodimer and rearrangement of subunit A12.2. eLife 8; 10.7554/eLife.43204 (2019).

56. Tegunov, D. & Cramer, P. Real-time cryo-electron microscopy data preprocessing with Warp. Nature methods 16, 1146–1152; 10.1038/s41592-019-0580-y (2019).

57. Zivanov, J. et al. RELION-3: new tools for automated high-resolution cryo-EM structure determination. bioRxiv; 10.1101/421123 (2018).

58. Webb, B. & Sali, A. Comparative Protein Structure Modeling Using MODELLER. Current protocols in protein science 86, 2.9.1-2.9.37; 10.1002/cpps.20 (2016).

59. Tunyasuvunakool, K. et al. Highly accurate protein structure prediction for the human proteome. Nature 596, 590–596; 10.1038/s41586-021-03828-1 (2021).

60. Fislage, M., Shkumatov, A. V., Stroobants, A. & Efremov, R. G. Assessing the JEOL CRYO ARM 300 for high-throughput automated single-particle cryo-EM in a multiuser environment. IUCrJ 7, 707–718; 10.1107/S2052252520006065 (2020).

61. Merk, A. et al. 1.8 Å resolution structure of β-galactosidase with a 200 kV CRYO ARM electron microscope. IUCrJ 7, 639–643; 10.1107/S2052252520006855 (2020).

62. Kato, T., Makino, F., Miyata, T., Horváth, P. & Namba, K. Structure of the native supercoiled flagellar hook as a universal joint. Nat Commun 10, 5295; 10.1038/s41467-019-13252-9 (2019).

63. Heiss, F. B., Daiß, J. L., Becker, P. & Engel, C. Conserved strategies of RNA polymerase I hibernation and activation. Nature communications 12, 758; 10.1038/s41467-021-21031-8 (2021).

64. Weaver, K. N. et al. Acrofacial Dysostosis, Cincinnati Type, a Mandibulofacial Dysostosis Syndrome with Limb Anomalies, Is Caused by POLR1A Dysfunction. The American Journal of Human Genetics 96, 765–774; 10.1016/j.ajhg.2015.03.011 (2015).

65. Ide, S., Imai, R., Ochi, H. & Maeshima, K. Transcriptional suppression of ribosomal DNA with phase separation. Science advances 6; 10.1126/sciadv.abb5953 (2020).

66. Schaefer, E. et al. Autosomal recessive POLR1D mutation with decrease of TCOF1 mRNA is responsible for Treacher Collins syndrome. Genet Med 16, 720–724; 10.1038/gim.2014.12 (2014).

67. Dauwerse, J. G. et al. Mutations in genes encoding subunits of RNA polymerases I and III cause Treacher Collins syndrome. Nat Genet 43, 20–22; 10.1038/ng.724 (2011).

68. Thiffault, I. et al. Recessive mutations in POLR1C cause a leukodystrophy by impairing biogenesis of RNA polymerase III. Nat Commun 6, 7623; 10.1038/ncomms8623 (2015).

69. Sanchez, E. et al. POLR1B and neural crest cell anomalies in Treacher Collins syndrome type 4. Genetics in medicine : official journal of the American College of Medical Genetics 22, 547–556; 10.1038/s41436-019-0669-9 (2020).

70. Gauquelin, L. et al. Clinical spectrum of POLR3-related leukodystrophy caused by biallelic POLR1C pathogenic variants. Neurology Genetics 5, e369; 10.1212/NXG.0000000000000369 (2019).

71. Kara, B. et al. Severe neurodegenerative disease in brothers with homozygous mutation in POLR1A. European journal of human genetics : EJHG 25, 315–323; 10.1038/ejhg.2016.183 (2017).

72. Cramer, P. et al. Structure of eukaryotic RNA polymerases. Annual review of biophysics 37, 337–352; 10.1146/annurev.biophys.37.032807.130008 (2008).

73. Smid, A., Riva, M., Bouet, F., Sentenac, A. & Carles, C. The association of three subunits with yeast RNA polymerase is stabilized by A14. The Journal of biological chemistry 270, 13534–13540; 10.1074/jbc.270.22.13534 (1995).

74. Gadal, O. et al. A34.5, a nonessential component of yeast RNA polymerase I, cooperates with subunit A14 and DNA topoisomerase I to produce a functional rRNA synthesis machine. Molecular and cellular biology 17, 1787–1795; 10.1128/MCB.17.4.1787 (1997).

75. Hayles, J. et al. A genome-wide resource of cell cycle and cell shape genes of fission yeast. Open biology 3, 130053; 10.1098/rsob.130053 (2013).

76. Darrière, T. et al. Genetic analyses led to the discovery of a super-active mutant of the RNA polymerase I. PLoS genetics 15, e1008157; 10.1371/journal.pgen.1008157 (2019).

77. Beckouet, F. et al. Two RNA polymerase I subunits control the binding and release of Rrn3 during transcription. Molecular and cellular biology 28, 1596–1605; 10.1128/MCB.01464-07 (2008).

78. Plaschka, C. et al. Architecture of the RNA polymerase II-Mediator core initiation complex. Nature 518, 376–380; 10.1038/nature14229 (2015).

79. Zimmermann, L. et al. A Completely Reimplemented MPI Bioinformatics Toolkit with a New HHpred Server at its Core. Journal of molecular biology 430, 2237–2243; 10.1016/j.jmb.2017.12.007 (2018).

80. Rong, H. et al. Structure of human upstream binding factor HMG box 5 and site for binding of the cell-cycle regulatory factor TAF1. Acta crystallographica. Section D, Biological crystallography 63, 730–737; 10.1107/S0907444907017027 (2007).

81. Stros, M., Launholt, D. & Grasser, K. D. The HMG-box: a versatile protein domain occurring in a wide variety of DNA-binding proteins. Cellular and molecular life sciences : CMLS 64, 2590–2606; 10.1007/s00018-007-7162-3 (2007).

82. Cary, P. D., Turner, C. H., Mayes, E. & Crane-Robinson, C. Conformation and domain structure of the non-histone chromosomal proteins, HMG 1 and 2. Isolation of two folded fragments from HMG 1 and 2. European journal of biochemistry 131, 367–374; 10.1111/j.1432-1033.1983.tb07272.x (1983).

83. Stros, M., Bacíková, A., Polanská, E., Stokrová, J. & Strauss, F. HMGB1 interacts with human topoisomerase IIalpha and stimulates its catalytic activity. Nucleic acids research 35, 5001–5013; 10.1093/nar/gkm525 (2007).

84. Panova, T. B., Panov, K. I., Russell, J. & Zomerdijk, J. C. B. M. Casein kinase 2 associates with initiation-competent RNA polymerase I and has multiple roles in ribosomal DNA transcription. Molecular and cellular biology 26, 5957–5968; 10.1128/MCB.00673-06 (2006).

85. Ray, S. et al. Topoisomerase IIα promotes activation of RNA polymerase I transcription by facilitating pre-initiation complex formation. Nature communications 4, 1598; 10.1038/ncomms2599 (2013).

86. Vanden Broeck, A. et al. Structural basis for allosteric regulation of Human Topoisomerase IIα. Nature communications 12, 2962; 10.1038/s41467-021-23136-6 (2021).

87. Canela, A. et al. Genome Organization Drives Chromosome Fragility. Cell 170, 507-521.e18; 10.1016/j.cell.2017.06.034 (2017).

88. Naidu, S., Friedrich, J. K., Russell, J. & Zomerdijk, J. C. B. M. TAF1B is a TFIIB-like component of the basal transcription machinery for RNA polymerase I. Science (New York, N.Y.) 333, 1640–1642; 10.1126/science.1207656 (2011).

89. Knutson, B. A. & Hahn, S. Yeast Rrn7 and human TAF1B are TFIIB-related RNA polymerase I general transcription factors. Science (New York, N.Y.) 333, 1637–1640; 10.1126/science.1207699 (2011).

90. Mars, J.-C., Sabourin-Felix, M., Tremblay, M. G. & Moss, T. A Deconvolution Protocol for ChIP-Seq Reveals Analogous Enhancer Structures on the Mouse and Human Ribosomal RNA Genes. G3 (Bethesda, Md.) 8, 303–314; 10.1534/g3.117.300225 (2018).

91. Cramer, P., Bushnell, D. A. & Kornberg, R. D. Structural basis of transcription: RNA polymerase II at 2.8 angstrom resolution. Science (New York, N.Y.) 292, 1863–1876; 10.1126/science.1059493 (2001).

92. Penrod, Y., Rothblum, K. & Rothblum, L. I. Characterization of the interactions of mammalian RNA polymerase I associated proteins PAF53 and PAF49. Biochemistry 51, 6519–6526; 10.1021/bi300408q (2012).

93. Knutson, B. A., McNamar, R. & Rothblum, L. I. Dynamics of the RNA polymerase I TFIIF/TFIIE-like subcomplex: a mini-review. Biochemical Society transactions 48, 1917–1927; 10.1042/BST20190848 (2020).

94. McNamar, R., Rothblum, K. & Rothblum, L. I. The Mammalian and Yeast A49 and A34 Heterodimers: Homologous but Not the Same. Genes 12; 10.3390/genes12050620 (2021).

95. Zomerdijk, J. Structural biology: Pivotal findings for a transcription machine. Nature 502, 629–630; 10.1038/nature12700 (2013).

96. Sadian, Y. et al. Molecular insight into RNA polymerase I promoter recognition and promoter melting. Nature communications 10, 5543; 10.1038/s41467-019-13510-w (2019).

97. Pilsl, M. & Engel, C. Structural basis of RNA polymerase I pre-initiation complex formation and promoter melting. Nature communications 11, 1206; 10.1038/s41467-020-15052-y (2020).

98. Cavanaugh, A. H. et al. Rrn3 phosphorylation is a regulatory checkpoint for ribosome biogenesis. The Journal of biological chemistry 277, 27423–27432; 10.1074/jbc.M201232200 (2002).

99. Schilbach, S., Aibara, S., Dienemann, C., Grabbe, F. & Cramer, P. Structure of RNA polymerase II pre-initiation complex at 2.9 Å defines initial DNA opening. Cell 184, 4064-4072.e28; 10.1016/j.cell.2021.05.012 (2021).

100. Aibara, S., Schilbach, S. & Cramer, P. Structures of mammalian RNA polymerase II pre-initiation complexes. Nature 594, 124–128; 10.1038/s41586-021-03554-8 (2021).

101. Abdella, R. et al. Structure of the human Mediator-bound transcription preinitiation complex. Science 372, 52–56; 10.1126/science.abg3074 (2021).

102. Liu, L. F. & Wang, J. C. Supercoiling of the DNA template during transcription. Proceedings of the National Academy of Sciences of the United States of America 84, 7024–7027; 10.1073/pnas.84.20.7024 (1987).

103. French, S. L. et al. Distinguishing the roles of Topoisomerases I and II in relief of transcription-induced torsional stress in yeast rRNA genes. Molecular and cellular biology 31, 482–494; 10.1128/MCB.00589-10 (2011).

104. Albert, B. et al. RNA polymerase I-specific subunits promote polymerase clustering to enhance the rRNA gene transcription cycle. The Journal of cell biology 192, 277–293; 10.1083/jcb.201006040 (2011).

105. Birch, J. L. et al. FACT facilitates chromatin transcription by RNA polymerases I and III. The EMBO journal 28, 854–865; 10.1038/emboj.2009.33 (2009).

106. Putnam, C. D., Copenhaver, G. P., Denton, M. L. & Pikaard, C. S. The RNA polymerase I transactivator upstream binding factor requires its dimerization domain and high-mobility-group (HMG) box 1 to bend, wrap, and positively supercoil enhancer DNA. Molecular and cellular biology 14, 6476–6488 (1994).

107. Morotomi-Yano, K. & Yano, K.-I. Nucleolar translocation of human DNA topoisomerase II by ATP depletion and its disruption by the RNA polymerase I inhibitor BMH-21. Sci Rep 11, 21533; 10.1038/s41598-021-00958-4 (2021).

108. Vos, S. M., Farnung, L., Linden, A., Urlaub, H. & Cramer, P. Structure of complete Pol II-DSIF-PAF-SPT6 transcription complex reveals RTF1 allosteric activation. Nat Struct Mol Biol 27, 668–677; 10.1038/s41594-020-0437-1 (2020).

109. Zhang, Y., Sikes, M. L., Beyer, A. L. & Schneider, D. A. The Paf1 complex is required for efficient transcription elongation by RNA polymerase I. Proceedings of the National Academy of Sciences 106, 2153–2158; 10.1073/pnas.0812939106 (2009).

110. Dodonova, S. O., Zhu, F., Dienemann, C., Taipale, J. & Cramer, P. Nucleosome-bound SOX2 and SOX11 structures elucidate pioneer factor function. Nature 580, 669–672; 10.1038/s41586-020-2195-y (2020).

111. McCullough, L. L. et al. Functional roles of the DNA-binding HMGB domain in the histone chaperone FACT in nucleosome reorganization. The Journal of biological chemistry 293, 6121–6133; 10.1074/jbc.RA117.000199 (2018).

112. Zhao, D. et al. Structure of the human RNA polymerase I elongation complex. Cell discovery 7, 97; 10.1038/s41421-021-00335-5 (2021).

113. Misiaszek, A. D. et al. Cryo-EM structures of human RNA polymerase I (2021).

114. Blum, M. et al. The InterPro protein families and domains database: 20 years on. Nucleic acids research 49, D344–D354; 10.1093/nar/gkaa977 (2021).

115. Katoh, K. & Standley, D. M. MAFFT multiple sequence alignment software version 7: improvements in performance and usability. Molecular Biology and Evolution 30, 772–780; 10.1093/molbev/mst010 (2013).

116. Castresana, J. Selection of conserved blocks from multiple alignments for their use in phylogenetic analysis. Molecular Biology and Evolution 17, 540–552; 10.1093/oxfordjournals.molbev.a026334 (2000).

117. Stamatakis, A. RAxML-VI-HPC: maximum likelihood-based phylogenetic analyses with thousands of taxa and mixed models. Bioinformatics (Oxford, England) 22, 2688–2690; 10.1093/bioinformatics/btl446 (2006).

118. Burki, F., Roger, A. J., Brown, M. W. & Simpson, A. G. B. The New Tree of Eukaryotes. Trends in Ecology & Evolution 35, 43–55; 10.1016/j.tree.2019.08.008 (2020).

119. Waterhouse, A. M., Procter, J. B., Martin, D. M. A., Clamp, M. & Barton, G. J. Jalview Version 2--a multiple sequence alignment editor and analysis workbench. Bioinformatics (Oxford, England) 25, 1189–1191; 10.1093/bioinformatics/btp033 (2009).

120. Gabler, F. et al. Protein Sequence Analysis Using the MPI Bioinformatics Toolkit. Current Protocols in Bioinformatics 72, e108; 10.1002/cpbi.108 (2020).

121. Panov, K. I., Friedrich, J. K., Russell, J. & Zomerdijk, J. C. B. M. UBF activates RNA polymerase I transcription by stimulating promoter escape. The EMBO journal 25, 3310–3322; 10.1038/sj.emboj.7601221 (2006).

122. Kastner, B. et al. GraFix: sample preparation for single-particle electron cryomicroscopy. Nature methods 5, 53–55; 10.1038/nmeth1139 (2008).

123. Palovcak, E. et al. A simple and robust procedure for preparing graphene-oxide cryo-EM grids. Journal of structural biology 204, 80–84; 10.1016/j.jsb.2018.07.007 (2018).

124. Schorb, M., Haberbosch, I., Hagen, W. J. H., Schwab, Y. & Mastronarde, D. N. Software tools for automated transmission electron microscopy. Nature methods 16, 471–477; 10.1038/s41592-019-0396-9 (2019).

125. Emsley, P. & Cowtan, K. Coot: model-building tools for molecular graphics. Acta crystallographica. Section D, Biological crystallography 60, 2126–2132; 10.1107/S0907444904019158 (2004).

126. Liebschner, D. et al. Macromolecular structure determination using X-rays, neutrons and electrons: recent developments in Phenix. Acta crystallographica. Section D, Structural biology 75, 861–877; 10.1107/S2059798319011471 (2019).

