## Supplementary Figures and Table for "The human RNA polymerase I structure reveals an HMG-like transcription factor docking domain specific to metazoans"

**Daiß et al.**

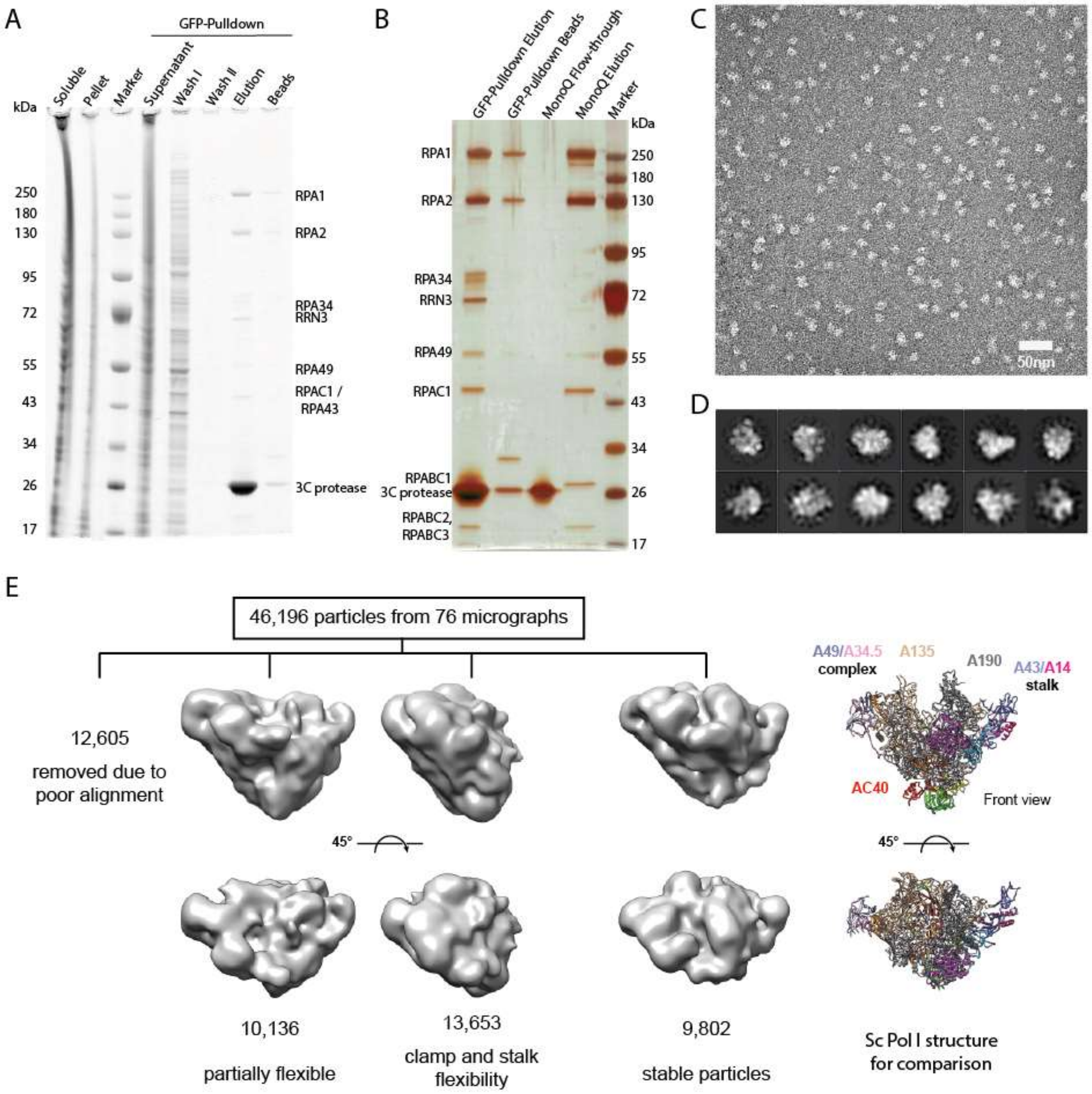

**Supplementary Figure 1. Human Pol I purification and negative stain EM**

**A** Coomassie-stained SDS-PAGE of human Pol I purification from lysates of the HeLa RPA1-sfGFP cell line. **B** Silver-stained SDS-PAGE shows loss of subunits RPA34, RPA49 and initiation factor RRN3 from most polymerases during MonoQ ion exchange chromatography. **C** Exemplary negative stain EM micrograph of hPol I eluted from anti-GFP-nanobody beads. **D** Exemplary 2D classes of picked particles. **E** Processing of negative stain EM data set by 3D classification shows flexible particles and ~21% of intact particles using this technique. Two orientations of each 3D class shown. Model of *S. cerevisiae* Pol I shown for comparison (right panel).

A

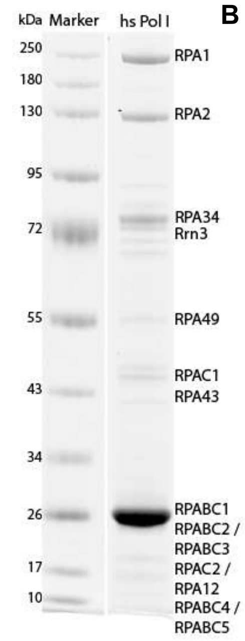

B

| subunit | score | No. of peptides | sequence coverage |
|---|---|---|---|
| RPA1 | 18730 | 233 | 75.7 % |
| RPA2 | 6765 | 106 | 58.9 % |
| RPAC1 | 3122 | 35 | 65.3 % |
| RPAC2 | 1604 | 22 | 66.2 % |
| RPA12 | 957 | 8 | 36.5 % |
| RPABC1 | 868 | 9 | 50.0 % |
| RPABC2 | 425 | 7 | 32.3 % |
| RPABC3 | 909 | 11 | 63.3 % |
| RPABC4 | 254 | 4 | 44.8 % |
| RPABC5 | 278 | 6 | 73.1 % |
| RPA43 | 564 | 9 | 29.0 % |
| RPA49 | 2525 | 39 | 53.2 % |
| RPA34 | 3770 | 50 | 71.2 % |
| RRN3 | 580 | 13 | 25.3 % |

C

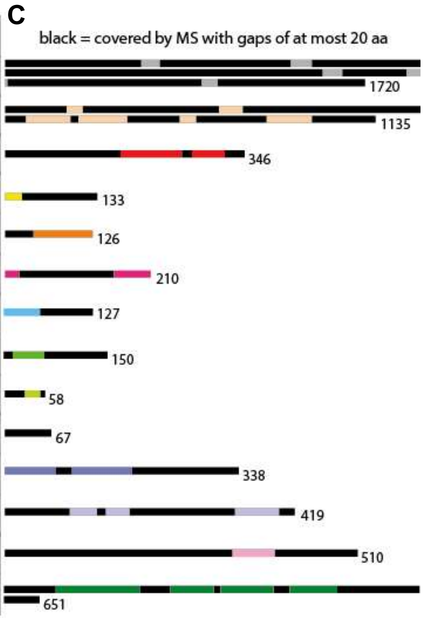

**Supplementary Figure 2. Mass Spectrometry analysis of human Pol I**

**A** Coomassie-stained SDS-PAGE of human Pol I purification; **B** MS-results for the subunits and initiation factor RRN3 labeled in Panel A with Sequence coverages of at least 25%. Number of identified peptides and score indicated; **C** Schematic representation of sequence coverage, to scale from N-terminus to C-terminus. Black bars indicate covered sequences (gaps <20 residues).

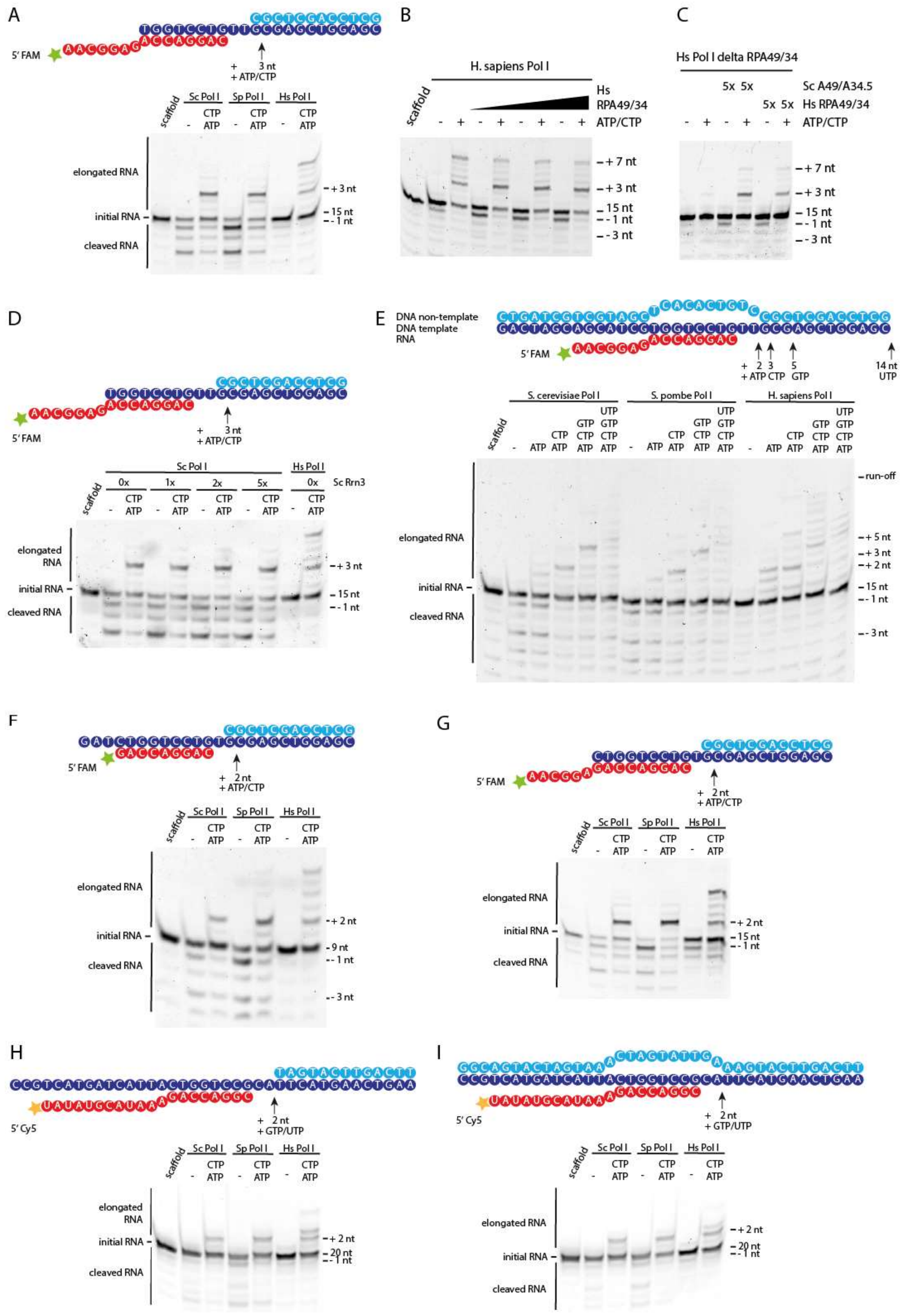

A
5′ FAM
+ 3 nt
+ ATP/CTP
scaffold
Sc Pol I
Sp Pol I
Hs Pol I
CTP ATP
elongated RNA
initial RNA
cleaved RNA
+ 3 nt
15 nt
- 1 nt
B
H. sapiens Pol I
Hs RPA49/34
ATP/CTP
+ 7 nt
+ 3 nt
15 nt
- 1 nt
- 3 nt
C
Hs Pol I delta RPA49/34
5x 5x
Sc A49/A34.5
Hs RPA49/34
ATP/CTP
+ 7 nt
+ 3 nt
15 nt
- 1 nt
- 3 nt
D
5′ FAM
+ 3 nt
+ ATP/CTP
Sc Pol I
Hs Pol I
0x 1x 2x 5x 0x
Sc Rrn3
elongated RNA
initial RNA
cleaved RNA
+ 3 nt
15 nt
- 1 nt
E
DNA non-template
DNA template
RNA
5′ FAM
+ 2 + ATP
3 CTP
5 GTP
14 nt UTP
S. cerevisiae Pol I
S. pombe Pol I
H. sapiens Pol I
ATP
CTP ATP
GTP CTP ATP
UTP GTP CTP ATP
run-off
+ 5 nt
+ 3 nt
+ 2 nt
15 nt
- 1 nt
- 3 nt
elongated RNA
initial RNA
cleaved RNA
F
5′ FAM
+ 2 nt
+ ATP/CTP
Sc Pol I
Sp Pol I
Hs Pol I
elongated RNA
initial RNA
cleaved RNA
+ 2 nt
9 nt
- 1 nt
- 3 nt
G
5′ FAM
+ 2 nt
+ ATP/CTP
Sc Pol I
Sp Pol I
Hs Pol I
elongated RNA
initial RNA
cleaved RNA
+ 2 nt
15 nt
- 1 nt
H
5′ Cy5
+ 2 nt
+ GTP/UTP
Sc Pol I
Sp Pol I
Hs Pol I
elongated RNA
initial RNA
cleaved RNA
+ 2 nt
20 nt
- 1 nt
I
5′ Cy5
+ 2 nt
+ GTP/UTP
Sc Pol I
Sp Pol I
Hs Pol I
elongated RNA
initial RNA
cleaved RNA
+ 2 nt
20 nt
- 1 nt

**Supplementary Figure 3. Proofreading ability of human Pol I is reduced compared to the yeast enzymes.**

Schematic Scaffold representations and Urea PAGE of elongation/cleavage assays show the activity of human Pol I compared to the *S. cerevisiae* and *S. pombe* homologues. **A** Basic scaffold shows differences in elongation / cleavage pattern between S. pombe, S. cerevisiae and H. sapiens enzymes. **B** Addition of recombinant human RPA49/34 does not abolish proofreading deficiency or induce deeper cleavage. **C** Human Pol I lacking the RPA49/34 heterodimer shows reduced activity. Complementation with recombinant human RPA49/34 or yeast A49/34.5 recovers elongation and cleavage activity but does not significantly change the pattern observed for complete hPol I (Panel A). Protein purification quality shown in Sup. Fig. 5. **D** Addition of the initiation factor Rrn3 does not impact elongation and cleavage functionality of Sc Pol I. **E - I** Experiments using different DNA/RNA scaffold sequences (top of each panel) show similar elongation patterns. Results from assays with scaffolds containing a complete nt-strand to create a mis-matched transcription bubble (**E** and **I**) underline the reduced backtracking ability of hPol I.

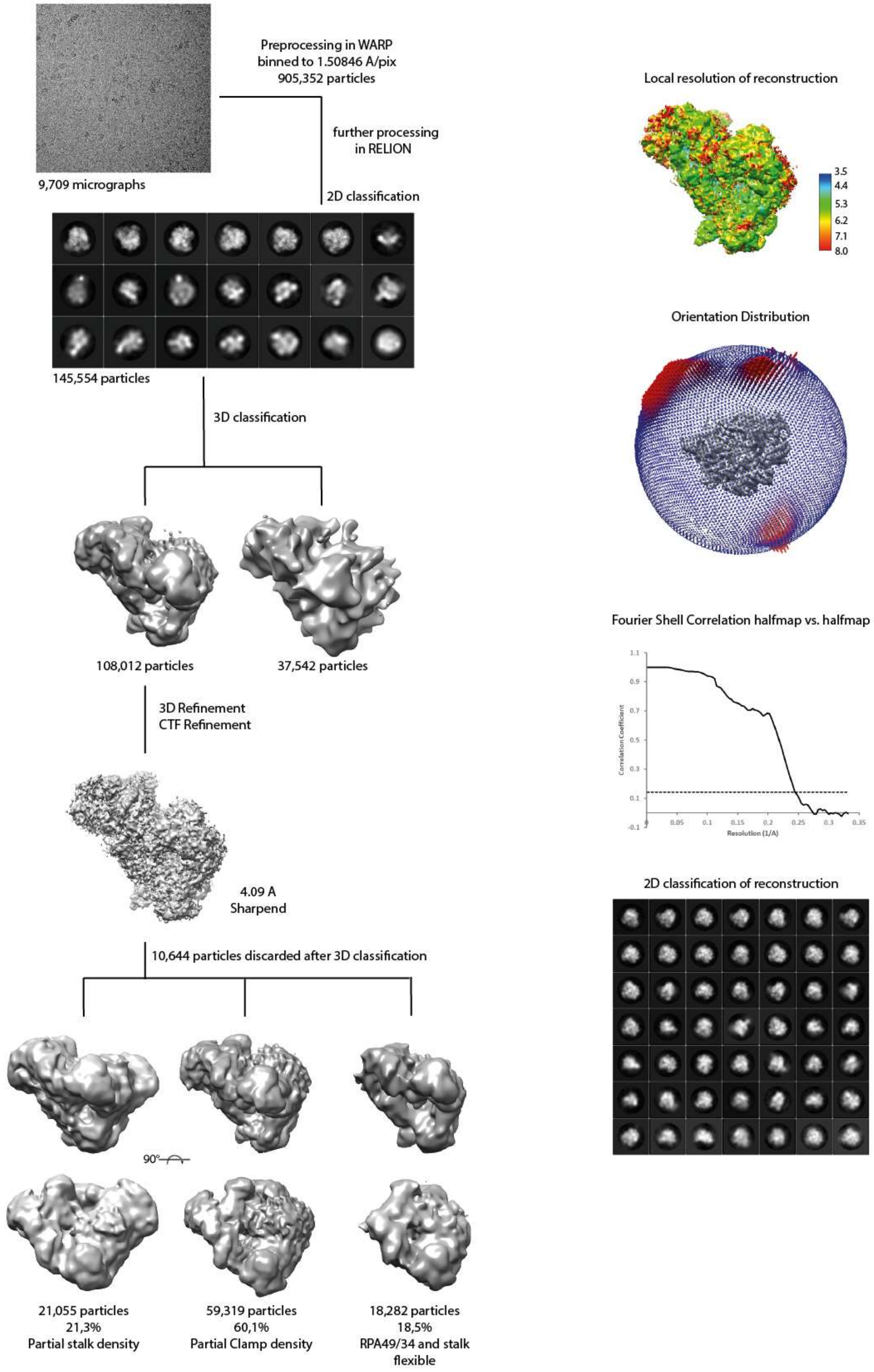

**Supplementary Figure 4. Human Pol I cryo-EM processing scheme**

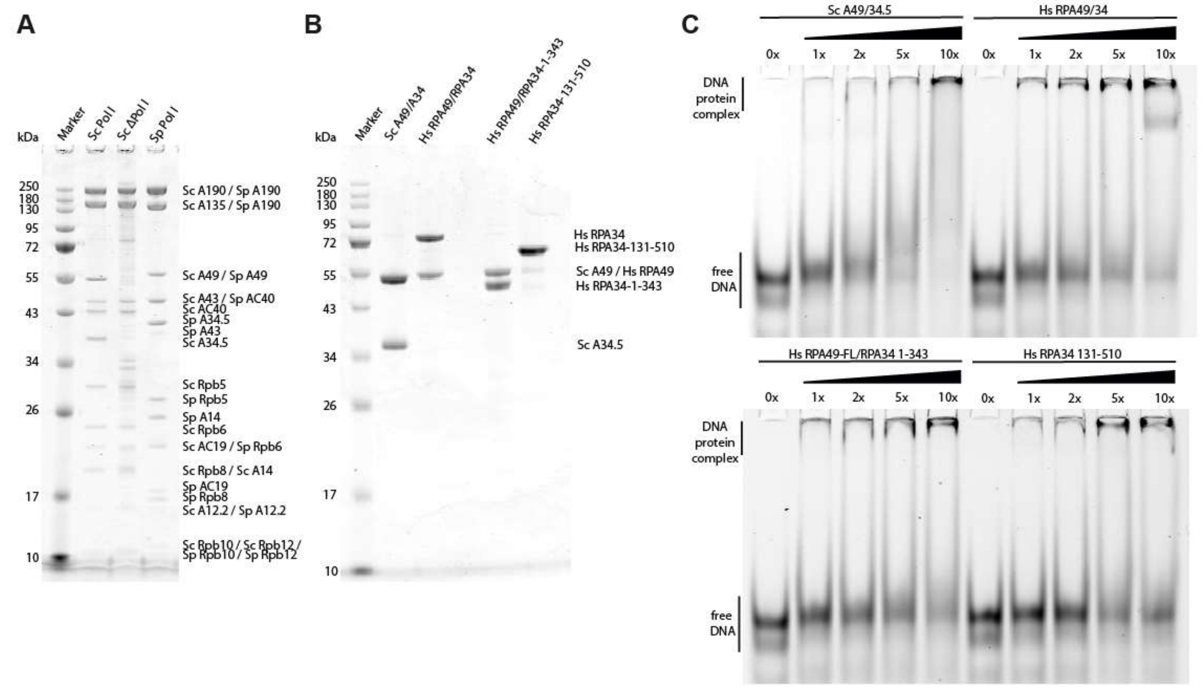

**Supplementary Figure 5. Protein purification and DNA-binding of RPA49/34**

**A** Coomassie-stained SDS-PAGE of purified yeast Pol I versions. **B** Coomassie-stained SDS-PAGE of purified *S. cerevisiae* and *H. sapiens* RPA49/34 versions. **C** EMSAs show binding of Hs RPA49/34 and its subdomains a 40bp dsDNA-fragment.

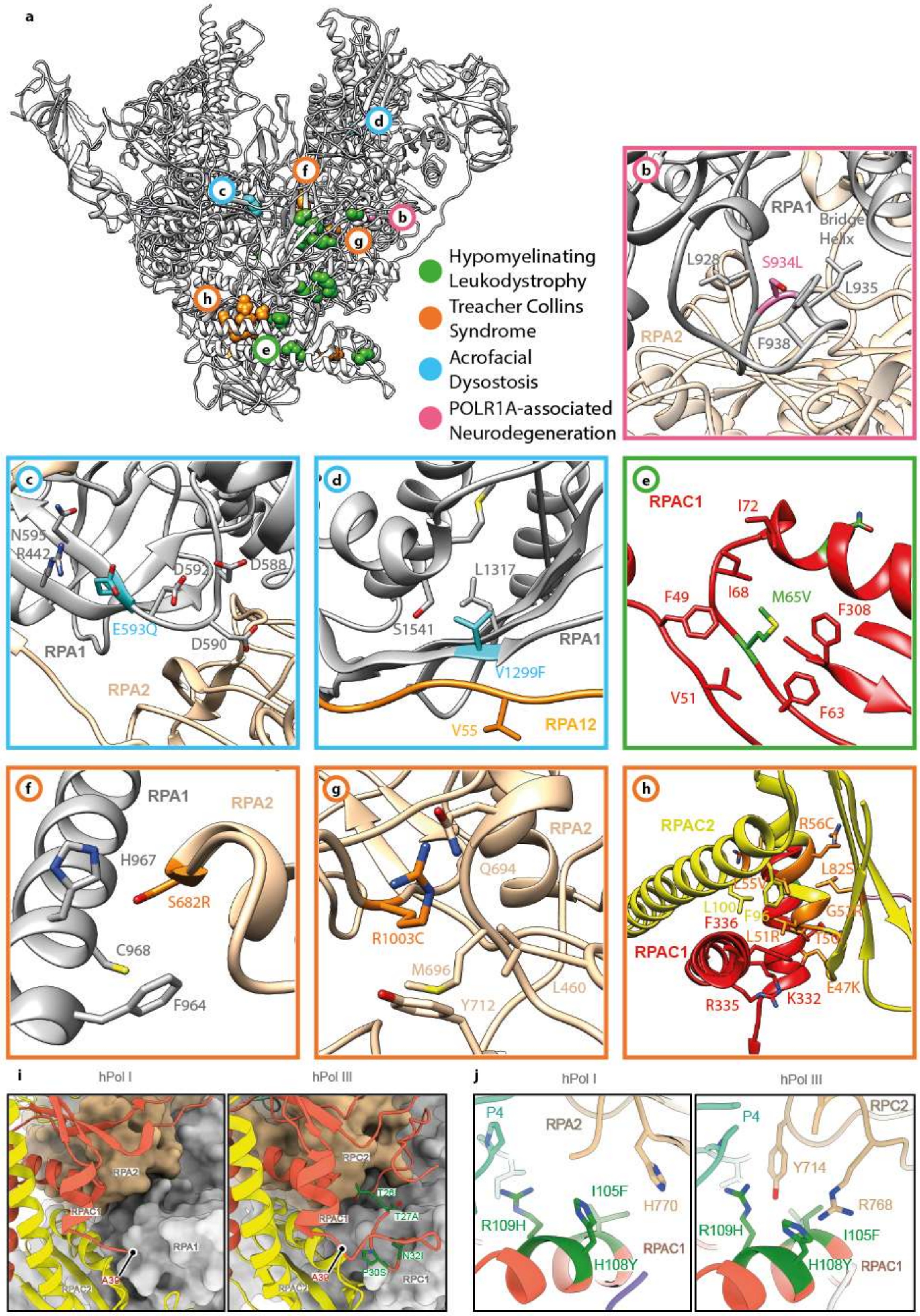

**Supplementary Figure 6. Location of hPol I residues mutated in disease**

**a** Structural model of hPol I (grey ribbon, back view) with location of disease-related mutation indicated (color code in panel). **b-h** Close-up views of the residues outlined in a. **i** The RPAC1 N-terminal region is flexible in hPol I compared to hPol III. **j** Interface of RPAC1 residues 105-109 with the second largest subunit in hPol I and hPol III.

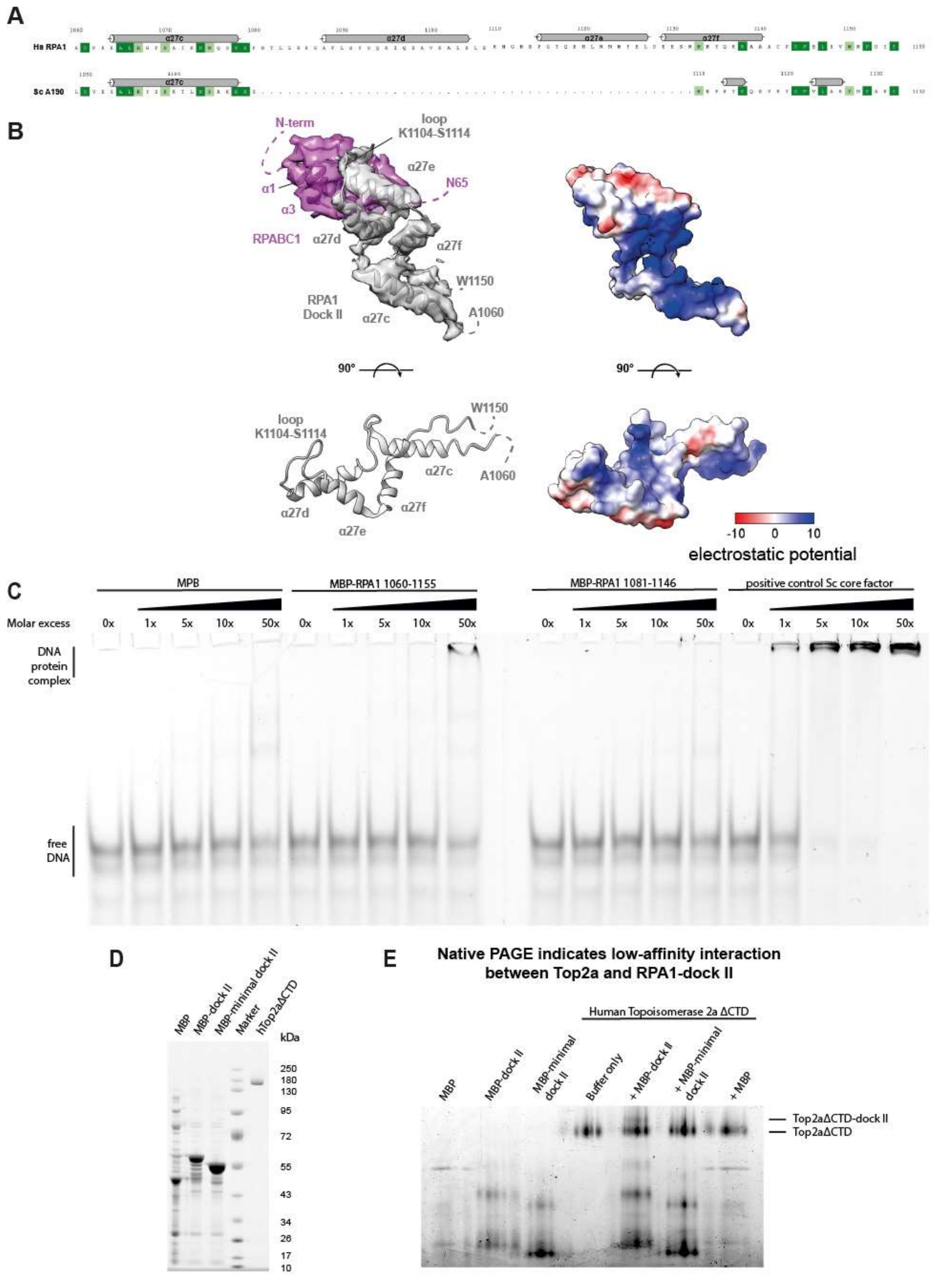

**Supplementary Figure 7. The HMG-box like dock II domain does not bind a dsDNA scaffold and transiently interacts with Top2a**

**A** Schematic representation of H.s. dock II in comparison to the yeast Pol I foot. **B** Electrostatic potential calculated in Chimera indicates an accessible basic patch. **C** EMSAs show low affinity of recombinant dock II to a 40bp dsDNA-fragment. Negative control: MBP-6xhis, positive control: S.c. ‘Core Factor’ **D** Coomassie-stained SDS-PAGE shows purified dock II-MBP fusion proteins and recombinant Top2a. **E** Native PAGE analysis shows a shift in the main Top2aΔCTD band in the presence of recombinant full-length MBP-dock II indicating a low-affinity interaction. The shift is not observed in a minimal dock II-MBP fusion or MBP only lanes.

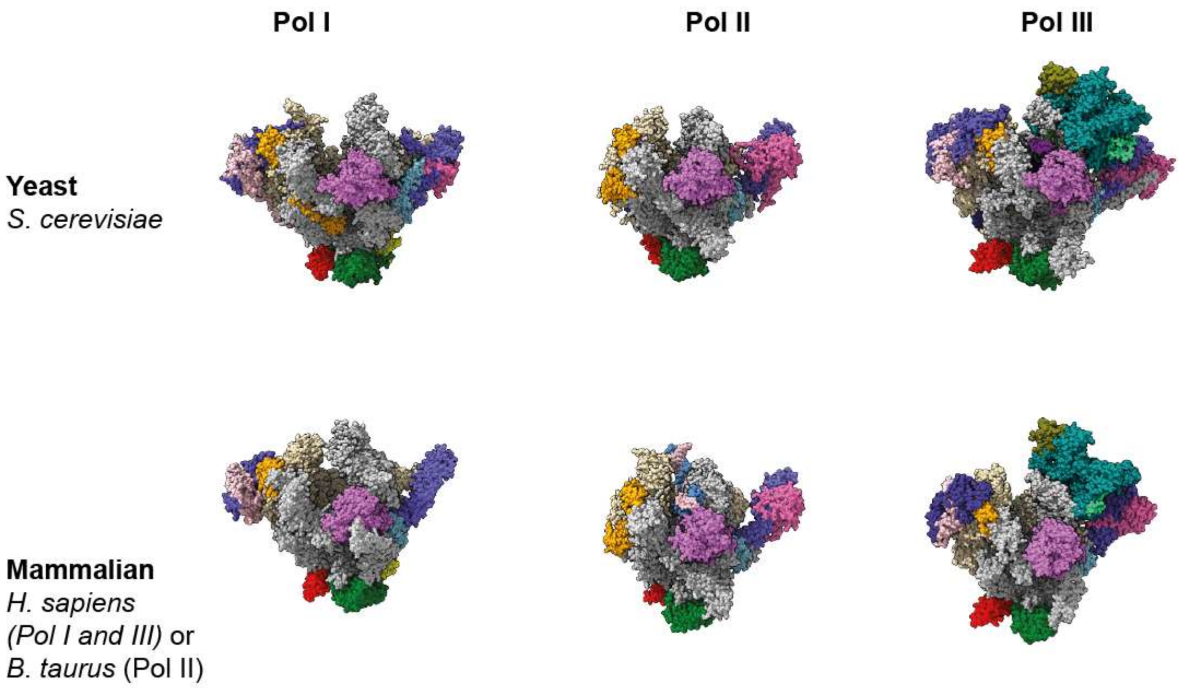

**Supplementary Figure 8. DNA-dependent RNA polymerase structures from yeast and mammalia**

Front view of Pol I, II and III from *S. cerevisiae* (top row; PDB 4C2M, 1WCM, 5FJ8) and from mammals (bottom row; this study, PDB 5FLM, 7AST). Subunits as in Table 1; Color code as in Fig. 2.

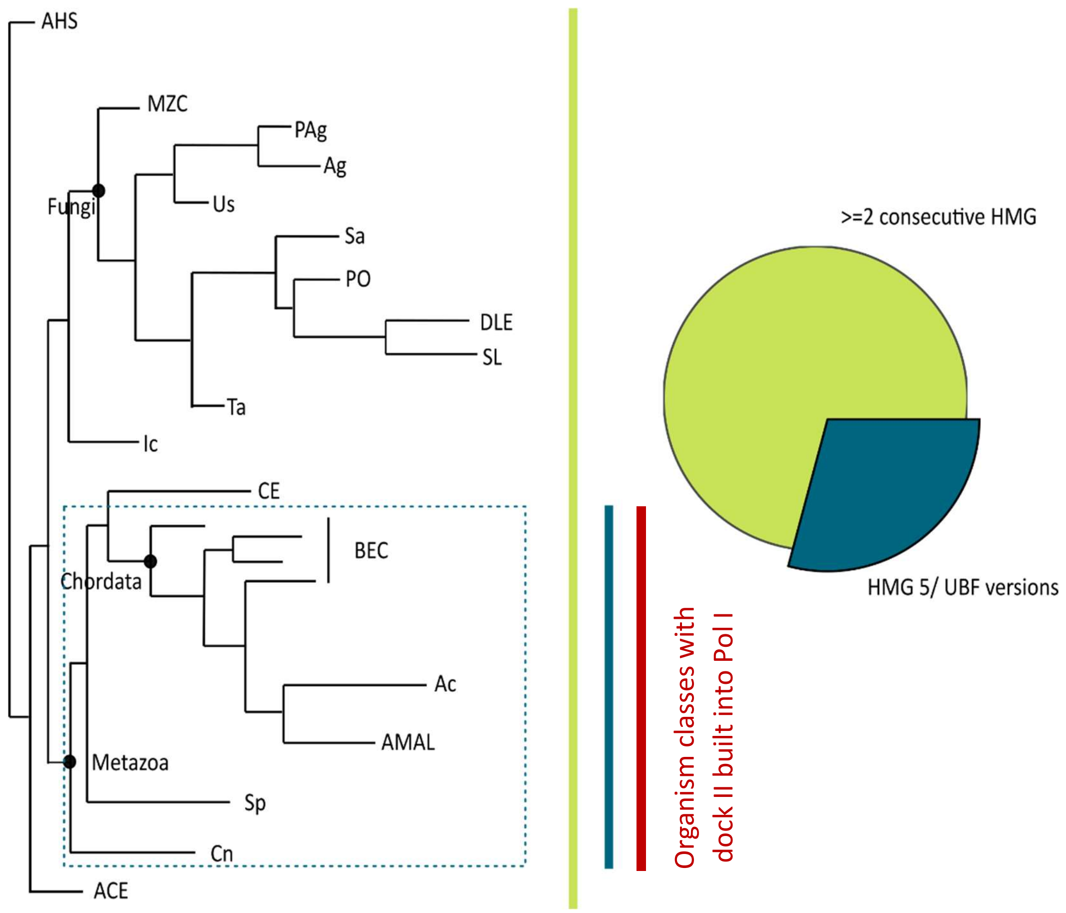

**Supplementary Figure 9. Taxonomic distribution of UBF versions**

Schematic phylogenetic tree based on Pol I subunit sequence homology (compare Fig. 3). HMG box domains (IPR009071) are present in HMO1 as also in UBF versions. As in both proteins a minimum of two consecutive boxes are present, the analyzed entries were restricted on such specific architecture. UBF-like proteins show five or six consecutive HMG boxes. Thus, the distribution of UBF proteins was restricted on the availability of the HMG box 5 (IPR029215). Taxonomic distribution of both entries was analyzed. A minimum of two consecutive HMG boxes can be found thoroughly over all organisms within our phylogenetic tree (green bar). The more specific HMG box 5 (UBF versions) is found in only 29% of the organisms (proportions shown in pie chart) clustering within the higher-order group of Metazoa (blue box and blue bar; Cryptophycea placed within Metazoa but belongs originally to the division of Cryptophyta). Ecdysozoa (within CE) seem to be more diverged from the group of Metazoa as only 1.5% of the sequences are annotated to have an HMG box 5. Red bar: Organisms classes in which the dock II domain was identified.

**Supplementary Table 1. Cryo-EM data collection and refinement statistics**

| | Human Pol I |
|---|---|
| **Data collection and processing** | |
| Magnification | 50.000 |
| Voltage (kV) | 200 |
| Electron exposure ($e^-$/Å$^2$) | 40 |
| Defocus range (μm) | -1.2 - -2.7 |
| Pixel size (Å) | 0.968 (binned to 1.5085) |
| Symmetry imposed | C1 |
| Initial particles images (no.) | 145,554 |
| Final particle images (no.) | 108,012 |
| Map resolution (Å) | 4.09 |
| FSC threshold | 0.143 |
| **Refinement** | |
| Initial model used (PDB code) | 5M3M |
| Model resolution (Å) | 3.5-4.1 Å |
| FSC threshold | 0.143 |
| Model composition | |
| Non-hydrogen atoms | 31,110 |
| Protein residues | 3,914 |
| Nucleotides | - |
| Ligands | - |
| B factors (Å$^2$) | |
| Protein | 212.67 |
| Nucleotides | - |
| Ligand | - |
| R.m.s. deviations | |
| Bond lengths (Å) | 0.008 |
| Bond angles (°) | 1.005 |
| Validation | |
| MolProbity score | 2.54 |
| Clashscore | 23.93 |
| Poor rotamers (%) | 0.41 |
| Ramachandran plot | |
| Favored (%) | 84.64 |
| Allowed (%) | 15.16 |
| Disallowed (%) | 0.21 |
