## Supplementary figures and images for "The human RNA polymerase I structure reveals an HMG-like transcription factor docking domain specific to metazoans"

### Supplementary Data 1

#### RPA1

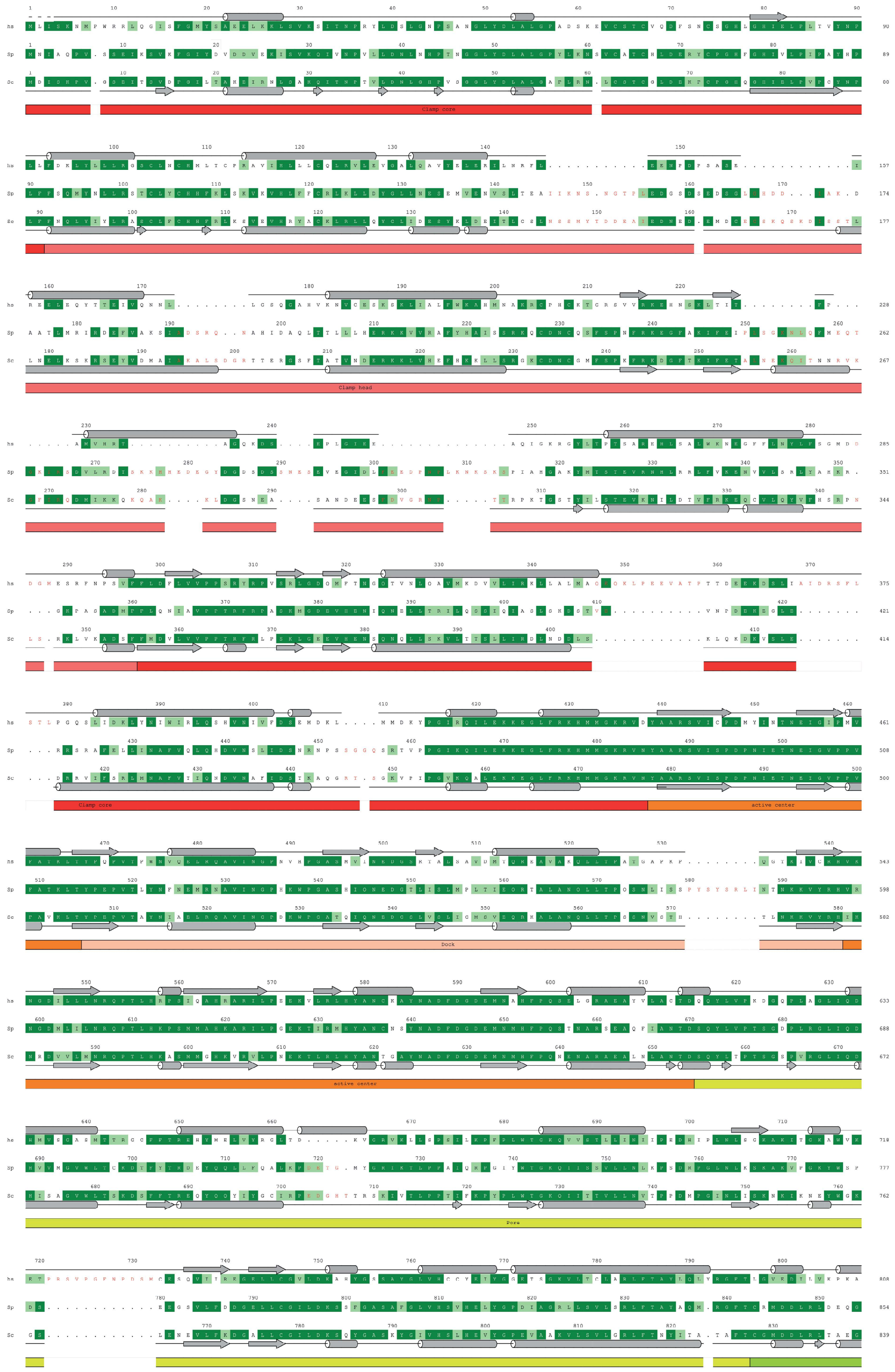

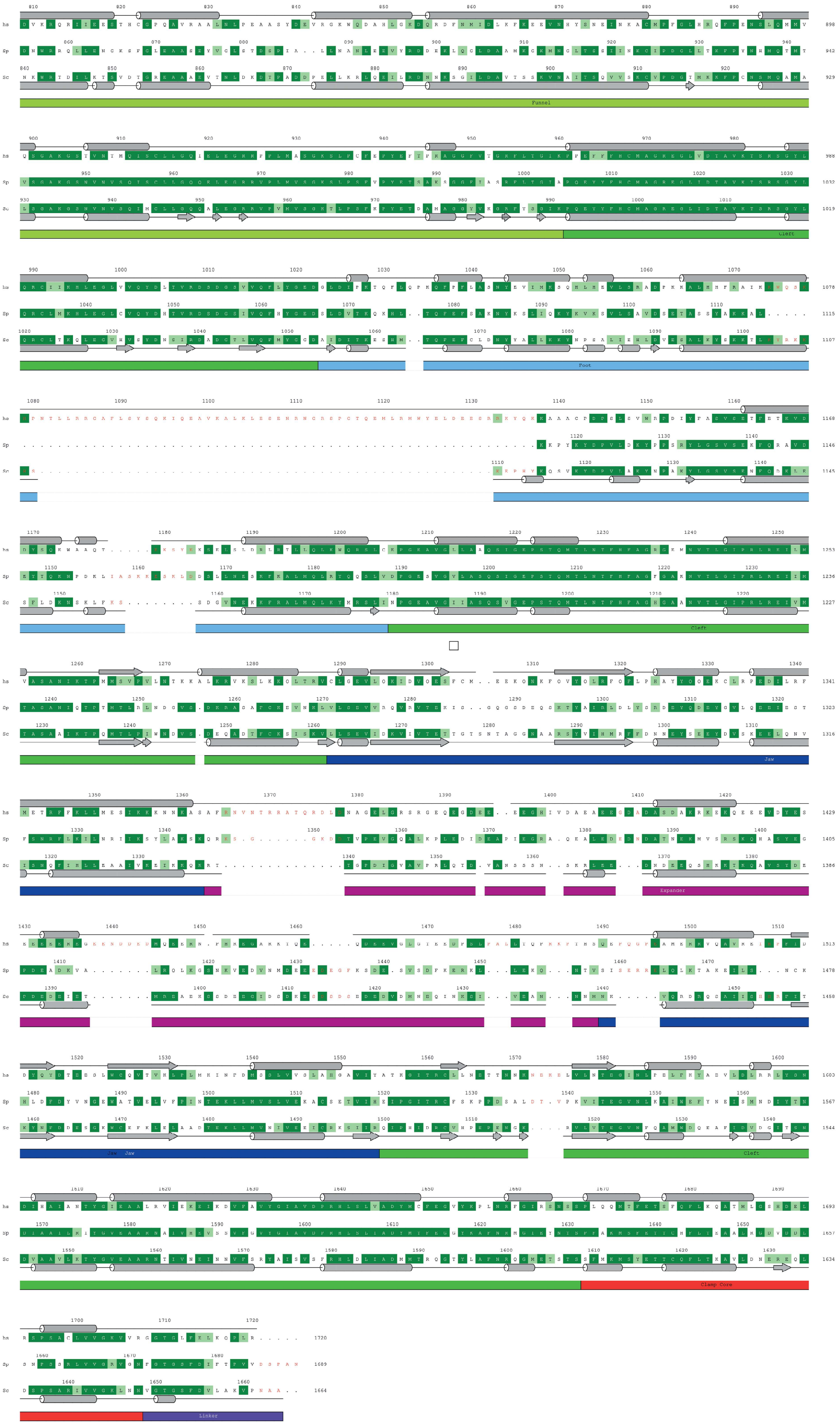

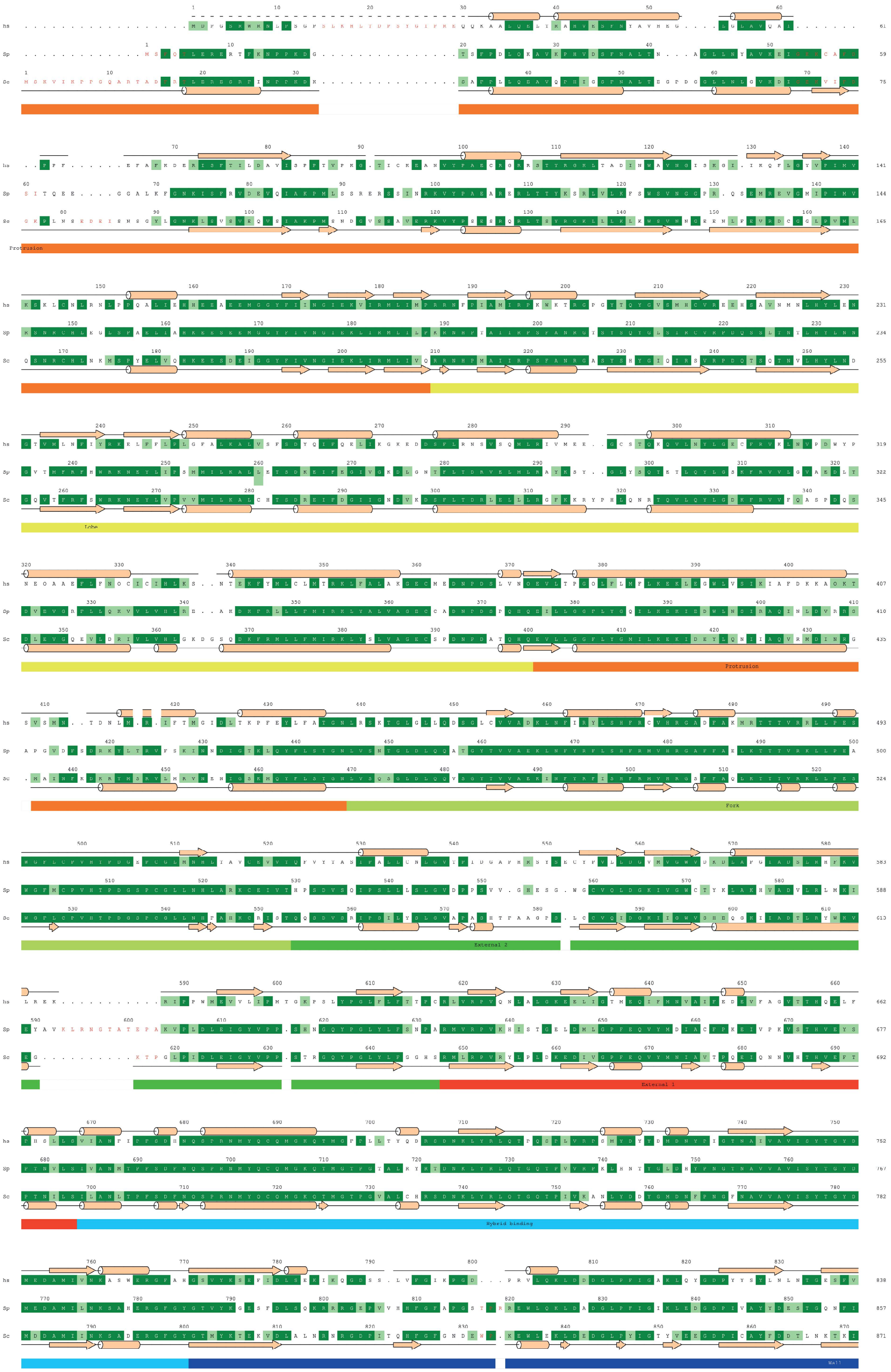

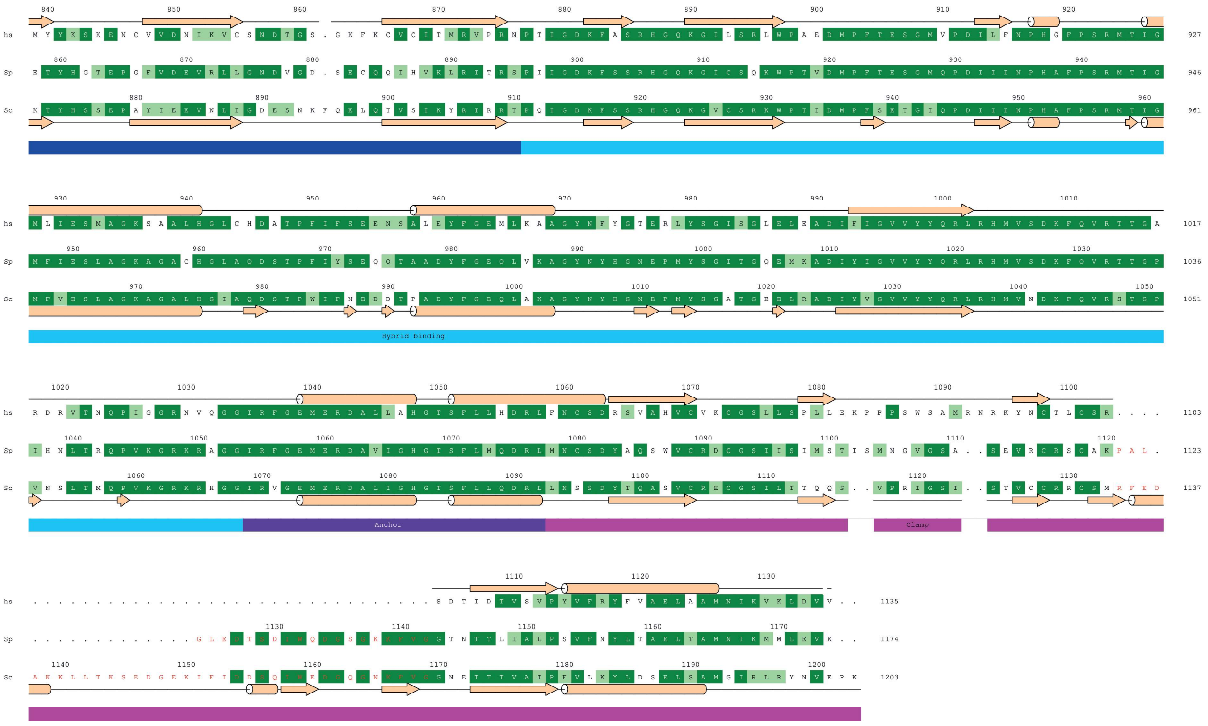

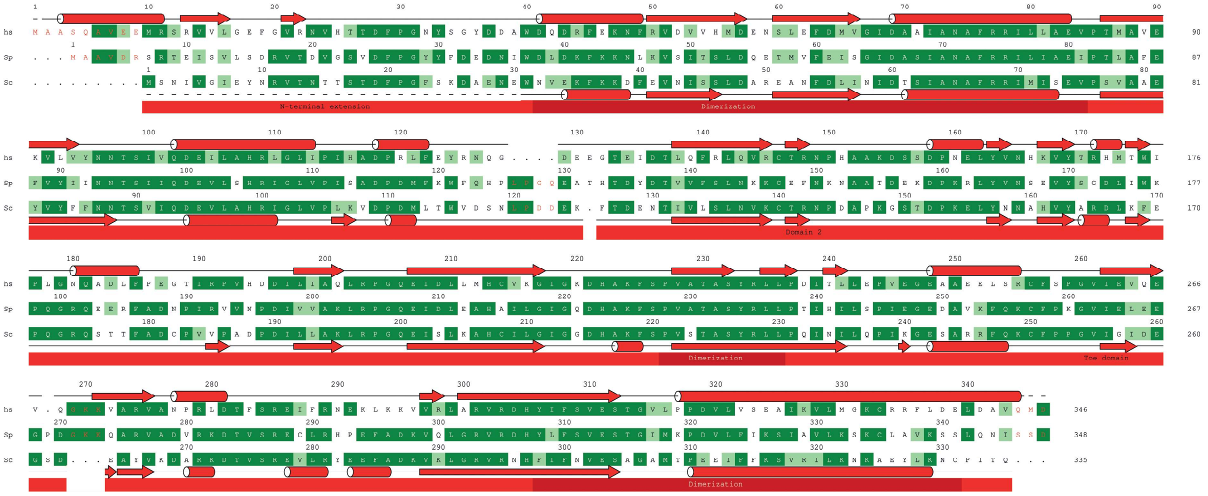

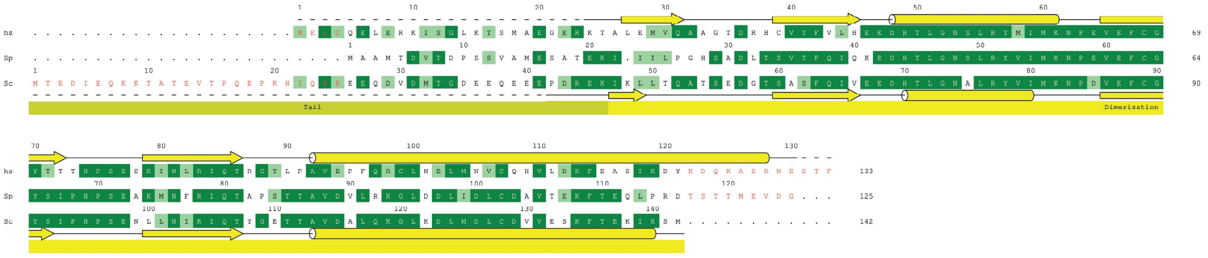

##### RPA49

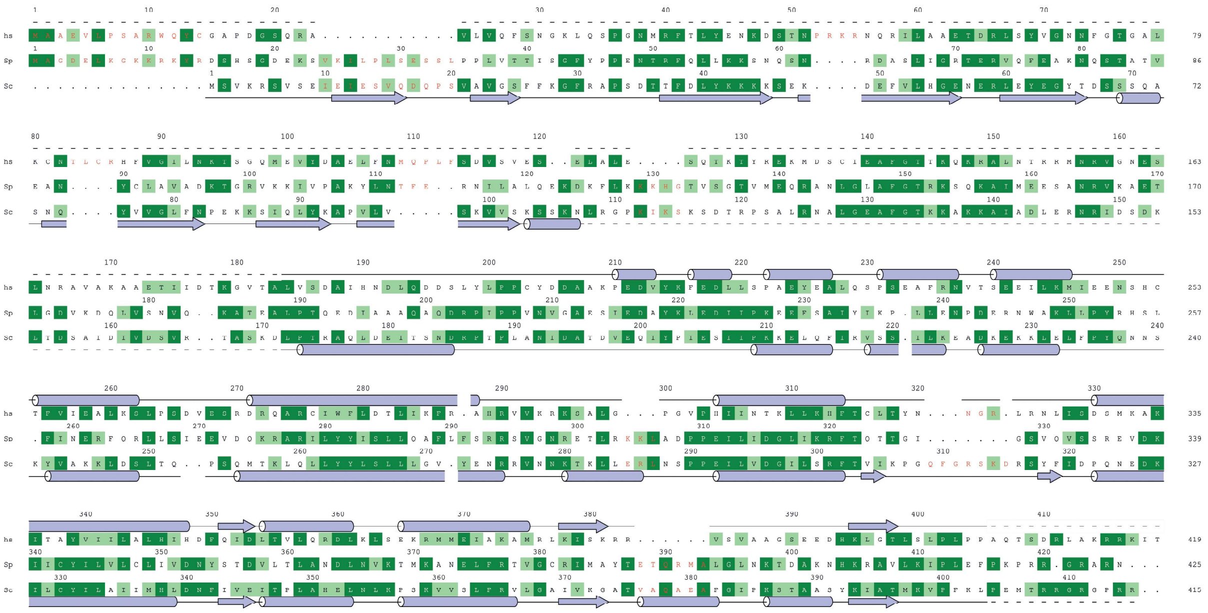


##### RPA34

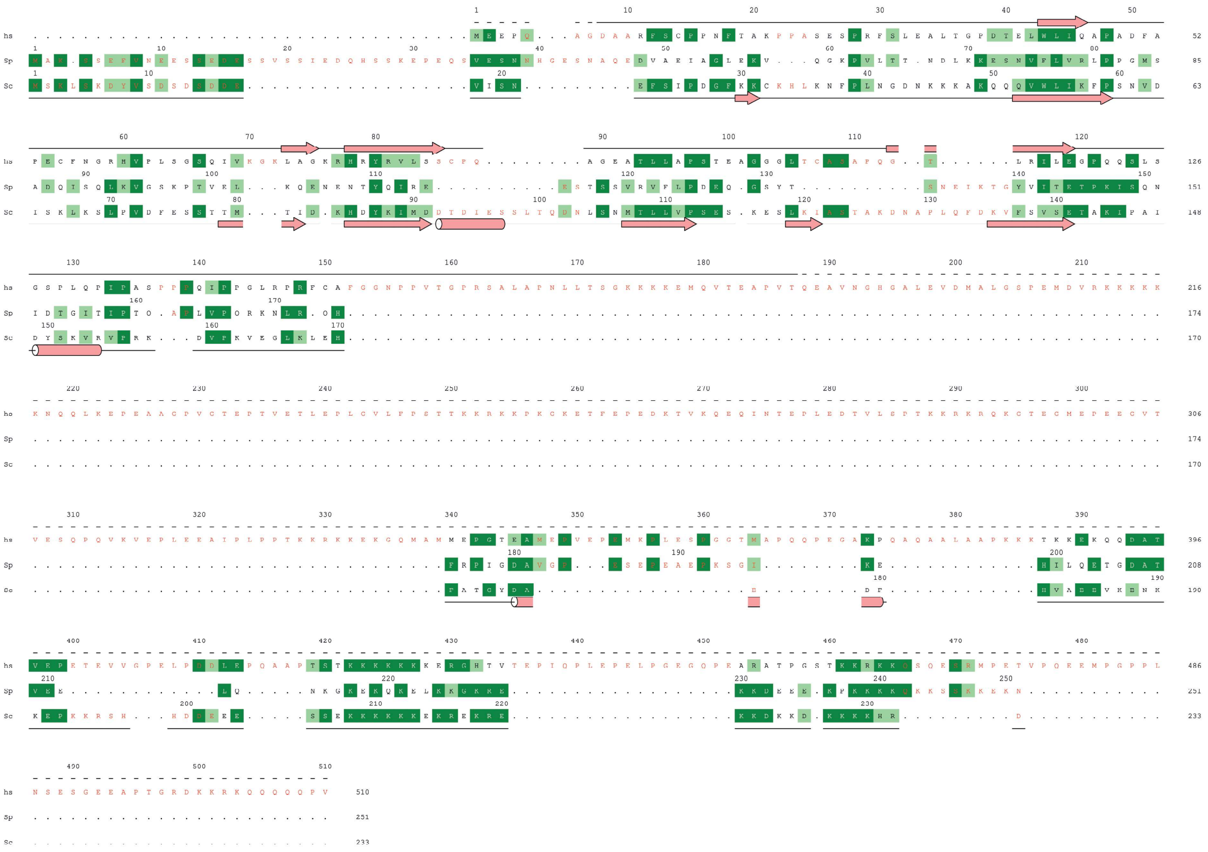


##### RPA12

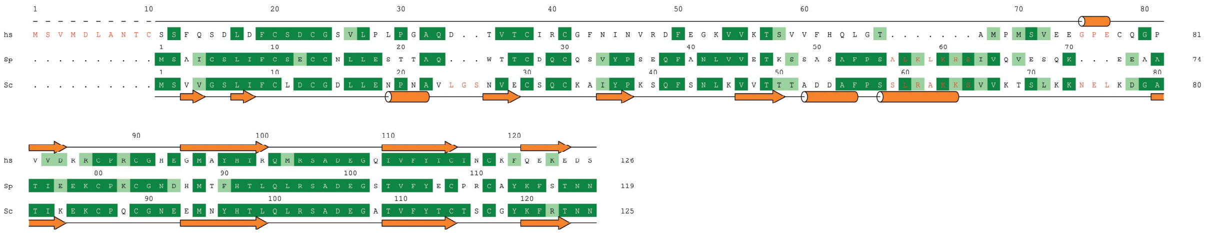

##### RPABC1

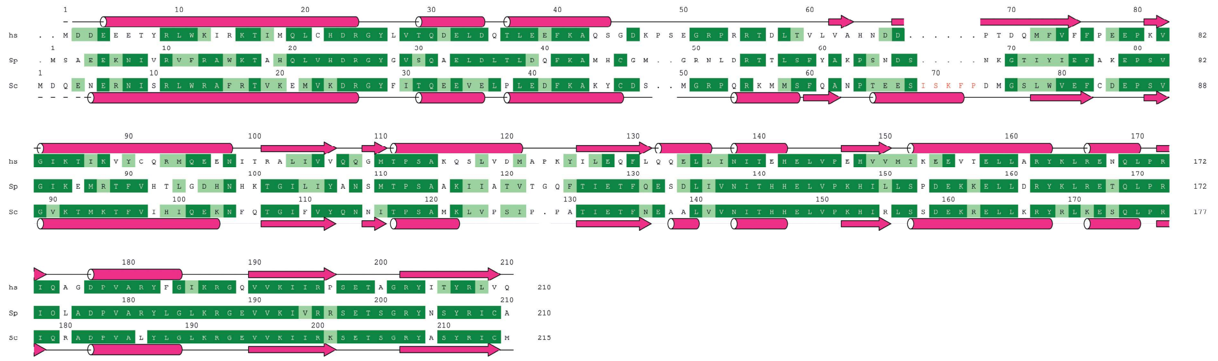

##### RPABC2

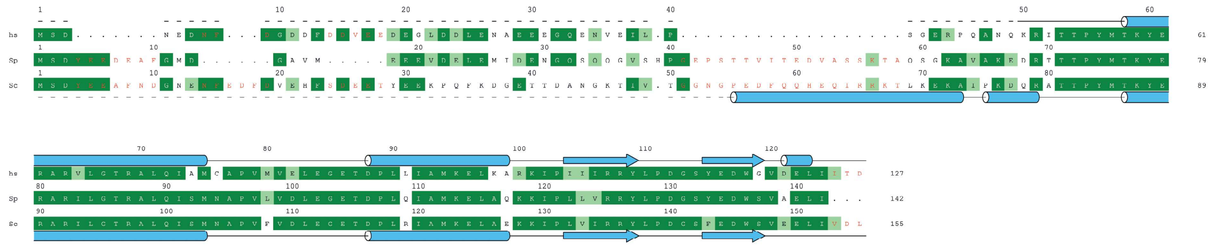

##### RPABC3

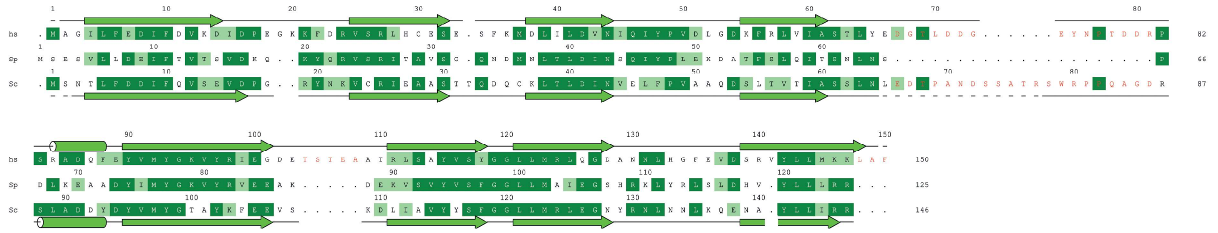

##### RPABC4

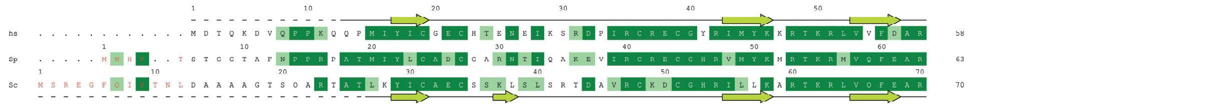

##### RPABC5

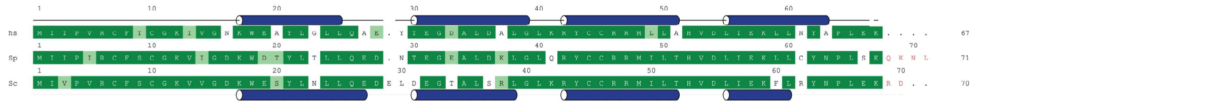

##### RPA43

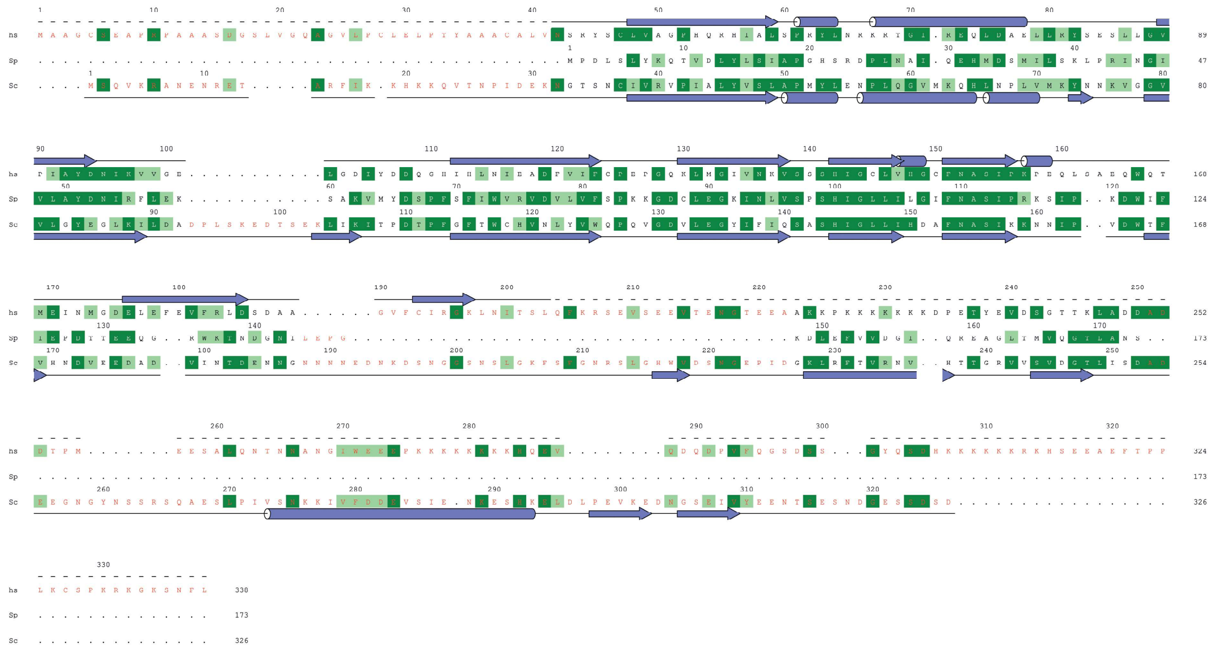
